# Minimization of proteome reallocation explains metabolic transition in hierarchical utilization of carbon sources

**DOI:** 10.1101/2024.01.23.576957

**Authors:** Zhihao Liu, Minghao Chen, Jingmin Hu, Yonghong Wang, Yu Chen

**Affiliations:** State Key Laboratory of Bioreactor Engineering, East China University of Science and Technology, Shanghai 200237, China; Key Laboratory of Quantitative Synthetic Biology, Shenzhen Institute of Synthetic Biology, Shenzhen Institute of Advanced Technology, Chinese Academy of Sciences, Shenzhen 518055, China

**Keywords:** enzyme constraint, metabolic model, mixed carbon sources, metabolic transition

## Abstract

Cells choose between alternative pathways in metabolic networks under diverse environmental conditions, but the principles governing the choice are insufficiently understood, especially in response to dynamically changing conditions. Here we observed that a lactic acid bacterium *Bacillus coagulans* displayed homolactic fermentation on glucose or trehalose as the sole carbon source, but transitioned from homolactic to heterolactic fermentation during the hierarchical utilization of glucose and trehalose when growing on the mixture. We simulated the observation by dynamic minimization of reallocation of proteome (dMORP) using an enzyme-constrained genome-scale metabolic model of *B. coagulans*, which coincided with our omics data. Moreover, we evolved strains to co-utilize mixed carbon sources and repress the choice of heterolactic fermentation, and the dynamics after co-utilization of carbon sources could also be captured by dMORP. Altogether, the findings suggest that upon rapid environmental changes bacteria tend to minimize proteome reallocation and accordingly adjust metabolism, and dMORP would be useful in simulating and understanding cellular dynamics.

## Introduction

Cells have alternative pathways, i.e., redundancy, in metabolic networks to adapt to diverse environmental conditions. For instance, while many microbes and mammalian cells use respiration for ATP production, some of them also harbor fermentation as an alternative ATP-producing pathway, which enables not only survival under anaerobic conditions but also fast growth in the presence of oxygen, i.e., aerobic fermentation(1). The alternative pathways empower cells to choose between distinct metabolic strategies, but the principles governing the cellular choice are insufficiently understood(2).

Experimental and theoretical studies suggest that the choices of alternative pathways and metabolic strategies reflect tradeoffs, mostly between metabolic efficiency, e.g., yield, of pathways and cellular resources, e.g., proteome, invested for enzymes of the pathways(3–8). For instance, fermentation has a lower ATP yield but is more proteome efficient than respiration, and therefore favors fast-growing cells subject to the limited proteome resources(9, 10). These studies are predominantly focused on steady-state cultures, e.g., chemostats, of model microbes, and therefore the cellular choice might be a result of a relatively long-term adaption to the environment. However, much less is known whether and how cells can transition between alternative pathways under dynamically changing environments, which are ubiquitous in both natural and artificial biological systems(11, 12).

Here we studied a lactic acid bacterium *Bacillus coagulans*, which has alternative pathways for catabolizing glycolytic carbon sources, e.g., glucose, and generating energy, i.e., homolactic and heterolactic fermentation(13, 14). The former converts glucose to lactate via the Embden– Meyerhof–Parnas (EMP) pathway with the theoretical yield being one gram lactate per gram glucose, while the latter via the phosphoketolase pathway with the halved yield due to the production of acetate and ethanol(15, 16). We observed that *B. coagulans* displayed the homolactic fermentation on glucose or trehalose as the sole carbon source, but transitioned from homolactic to heterolactic fermentation in the hierarchical utilization of glucose and trehalose when growing on the mixture. We hypothesized that the metabolic transition could be interpreted by minimization of reallocation of proteome (MORP) and performed enzyme-constrained metabolic modeling and omics analysis to test the hypothesis. Finally, by adaptive laboratory evolution (ALE) we evolved strains that can co-utilize the mixed carbon sources, which showcased another application of MORP.

## Results

### Lactate yield significantly changed in hierarchical utilization of glucose and trehalose

*B. coagulans* is widely used in the industrial lactate fermentation(14, 17, 18), where hydrolysates of low-cost starch-based materials are commonly used as the carbon sources(19, 20) considering cost and benefit. However, the co-existence of multiple carbon sources in the medium may lead to the hierarchical utilization(21–26) and thus the waste of substrates(27), the extension of fermentation cycle (28) and the difficulty in product separation and purification(29). Notably, the hierarchical utilization with the transition from homolactic to heterolactic fermentation led to the significantly reduced yield of lactate in the industrial fermentation by *B. coagulans*(13, 14). Therefore, identifying the principles governing the dynamic behavior of *B. coagulans* during the hierarchical utilization of mixed carbon sources would be of significance from both basic and applied science perspectives.

We performed batch fermentation of *B. coagulans* on the mixture of glucose and trehalose, a disaccharide with a high percentage in the hydrolysates(30). We observed the hierarchical utilization of glucose and trehalose, which divided the entire fermentation process into three phases (Fig. 1A). In the glucose utilization phase (P_gluc_, 0-12 h), glucose was exclusively consumed and lactate was rapidly produced with the lactate yield on glucose being 0.897 g/g, which was similar to the yield (i.e., 0.882 g/g) on glucose as the sole carbon source (Fig. 1B) and close to the yield of homolactic fermentation(15, 31). Upon glucose depletion, the lag phase occurred (P_lag_, 12-16 h), in which the lactate production stopped, the biomass concentration (measured by OD_620_) declined (Fig. 1A) and a few organic acids were slightly consumed (Supplementary Fig. 1). The biomass decline was also observed in the growth profiles on a mixture of glucose and xylose and switchgrass hydrolysate(32). We confirmed that the growth arrest was not caused by the depletion of nutrients such as amino acids (Supplementary Fig. 2). In the trehalose utilization phase (P_tre_, 16- 84 h), trehalose was consumed, and lactate was produced again (Fig. 1A). Interestingly, the lactate yield on trehalose in P_tre_ was only 0.526 g/g, which was much lower than that (i.e., 0.851 g/g) on trehalose as the sole substrate (Fig. 1B) and close to the theoretical lactate yield of heterolactic fermentation(15). This was in line with the increased yields of byproducts in P_tre_ (Supplementary Table 1). The significant change in the lactate yield during the hierarchical utilization of glucose and trehalose indicated distinct strategies to catabolize the carbon sources in P_gluc_ and P_tre_.

**Figure 1.**
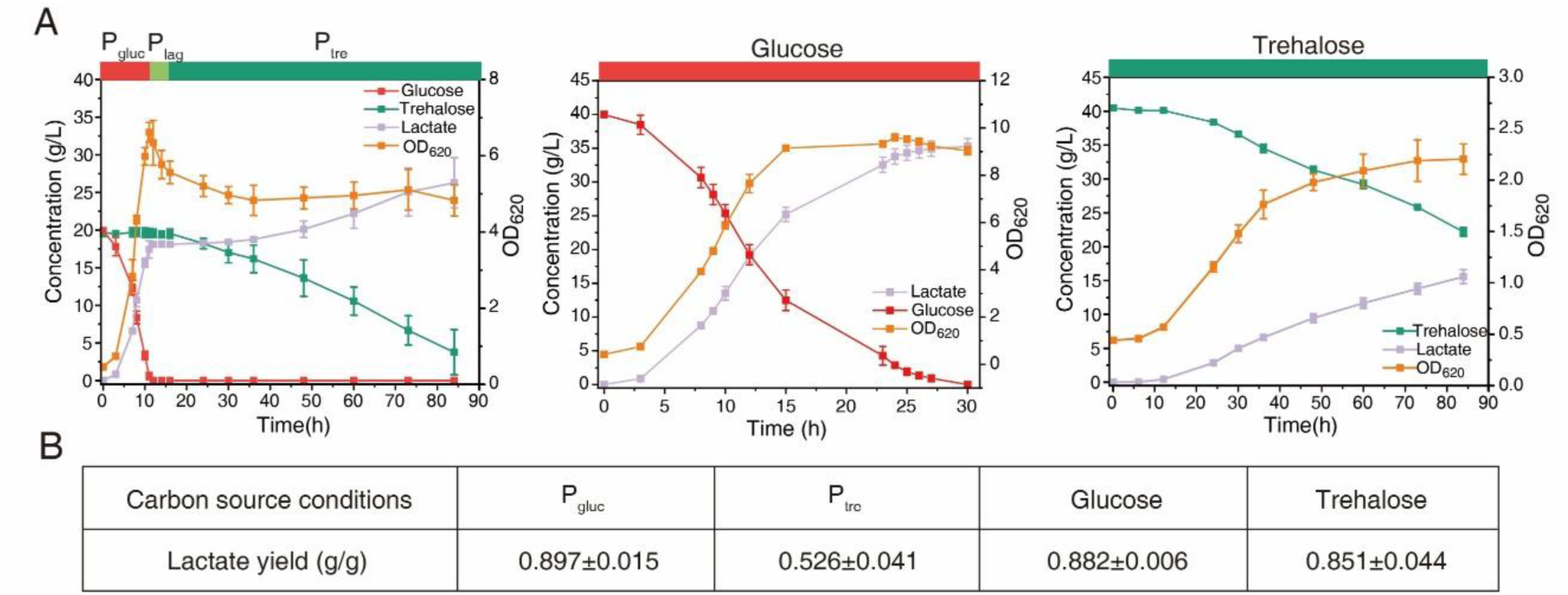
Fermentation on various carbon sources in 5 L bioreactor by *B. coagulans*. (A) Fermentation processes on glucose and trehalose as the mixed carbon sources (left), sole glucose (middle) and sole trehalose (right). Data in the plots are shown with mean ± s.d. of three biological replicates. (B) Lactate yields under various phases and conditions. Data in the table are shown with mean ± s.d. of three biological replicates.

### Dynamic minimization of reallocation of proteome predicted metabolic changes

To gain a systematic understanding, we reconstructed a genome-scale metabolic model of the strain *B. coagulans* DSM 1 = ATCC 7050(33) called iBcoa620, which was evaluated by the Memote score(34) and validated by growth on various carbon sources (Supplementary Fig. 3). Subsequently, we converted iBcoa620 into an enzyme-constrained version eciBcoa620 using the GECKO toolbox(35), allowing for understanding of metabolic changes by taking into account the enzyme usage and proteome allocation(36–38).

To predict the changes during the fermentation process, we performed dynamic flux balance analysis (dFBA)(39) with eciBcoa620 (Fig. 2A). The dFBA approach can divide the entire period into multiple time intervals and obtain the flux distribution at each time interval by performing classical FBA(40) which requires a predefined objective function. While growth maximization(41) as the objective function appeared reasonable to simulate for P_gluc_, it could not predict the cellular behavior after glucose depletion (Fig. 2B), meaning that cells might adjust the objective in response to the changing environment. Therefore, we attempted other commonly used objective functions(42), including to maximize lactate production and to maximize non-growth-associated maintenance (NGAM), and found that they also failed to capture the cellular behavior in P_tre_ (Supplementary Fig. 4), i.e., predicted either much higher lactate concentrations than measurements or no lactate production (Fig. 2B).

**Figure 2.**
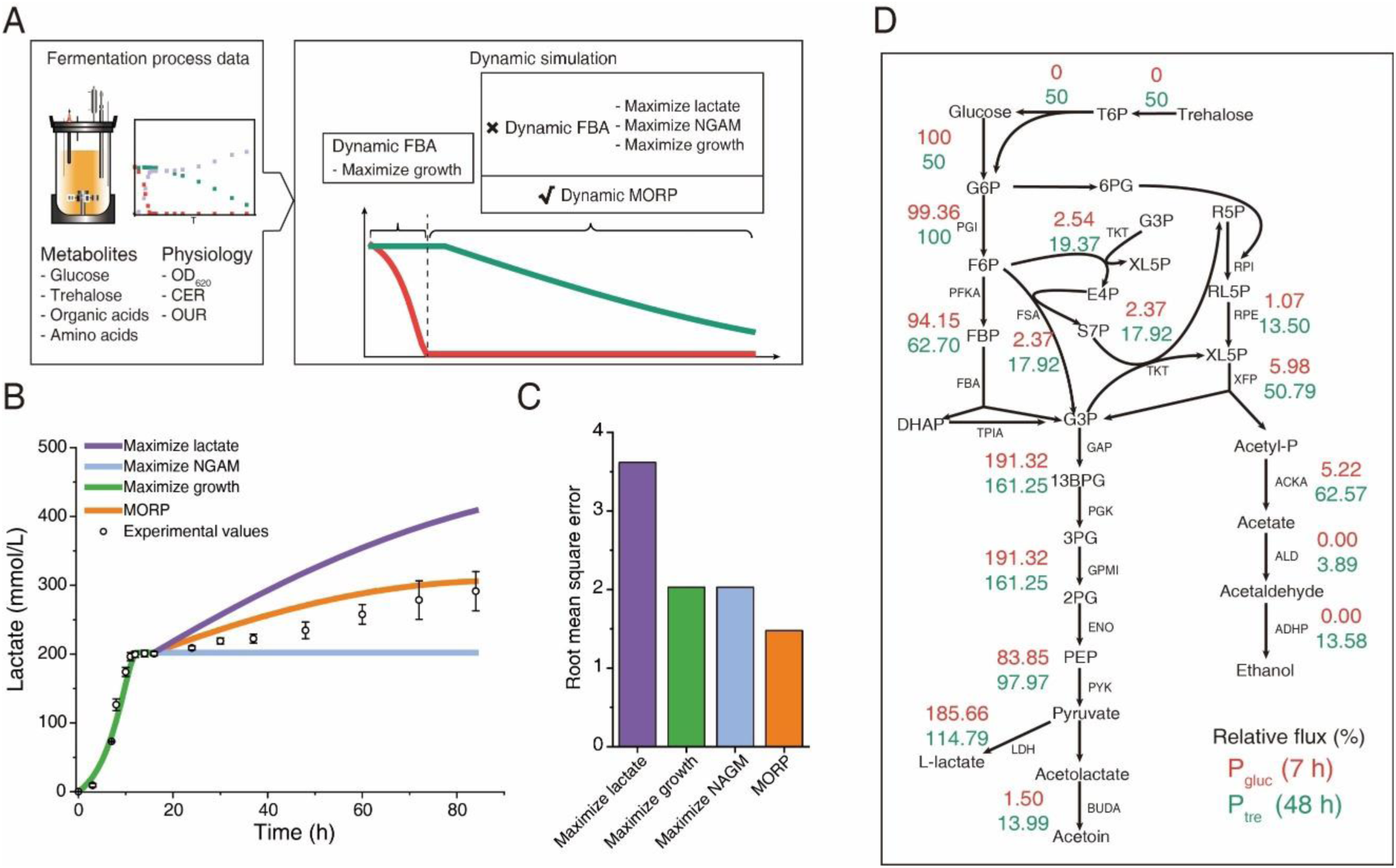
Simulation of batch fermentation under mixed carbon sources by eciBcoa620. (A) Simulation of the fermentation process. In the dFBA simulation of P_gluc_ the growth rate was maximized. To simulate P_tre_, various objective functions could be adopted within the dFBA framework. Additionally, we developed an approach to dynamically minimize the proteome reallocation with the enzyme-constrained models. (B) Simulation of lactate production using different objective functions. Minimization of proteome reallocation predicted the lactate production in P_tre_ better than the others including maximizing lactate, NGAM and growth. Data in the plot are shown with mean ± s.d. of three biological replicates. (C) Root mean square errors of the different objective functions in predicting P_tre_. Root mean square error quantifies the difference between the predicted and experimental concentrations of detected products, including lactate, acetate, pyruvate, acetoin and ethanol. (D) Comparison of the relative fluxes of the central carbon metabolism (CCM) between P_gluc_ (represented by 7 h) and P_tre_ (represented by 48h). Relative flux is calculated as the percentage of the absolute flux of each reaction to the substrate uptake flux. Note that one unit of trehalose uptake flux was converted to two units of glucose uptake flux. Abbreviations: T6P, trehalose 6-phosphate; G6P, glucose 6-phosphate; F6P, fructose 6 - phosphate; FBP, fructose 1,6 - bisphosphate; DHAP, dihydroxyacetone phosphate; G3P, glyceraldehyde 3-phosphate; 13BPG, glycerate 1,3-bisphosphate; 3PG, 3-phosphoglycerate; 2PG, 2-Phospho-D-glycerate; PEP, phosphoenolpyruvate; 6PG, gluconate 6-phosphate; E4P, erythrose 4-phosphate; S7P, sedoheptulose 7-phosphate; R5P, ribose 5-phosphate; RL5P, ribulose 5-phosphate; XL5P, xylulose 5-phosphate; Acetyl-P, Acetyl phosphate; PGI, glucose-6- phosphate isomerase; PKFA, 6-phosphofructokinase; FBA, fructose-1,6-bisphosphate aldolase; GAP, glyceraldehyde-3-phosphate dehydrogenase; PGK, phosphoglycerate kinase; GPMI, phosphoglycerate mutase; ENO, enolase; PYK, pyruvate kinase; LDH, L-lactate dehydrogenase; RPI, ribose 5-phosphate isomerase; RPE, ribulose-phosphate 3-epimerase; XFP, phosphoketolase; FSA, fructose-6-phosphate aldolase; TKT, transketolase; BUDA, acetolactate decarboxylase; ACKA, acetate kinase; ALD, aldehyde dehydrogenase; ADHP, alcohol dehydrogenase.

Considering the fact that proteome reallocation requires frequent protein synthesis and degradation, which is of high cost to cells(42), we hypothesized that cells tend to dynamically minimize the proteome reallocation in response to rapid environmental changes. To test it, we developed the approach MORP and extended to the dynamic version dMORP to simulate the cellular behavior after glucose depletion, and the enzyme usage in the last time point of the glucose phase was the reference for the first time point for of the simulations by dMORP (Fig. 2A) (Materials and Methods). The dMORP approach leverages the enzyme-constrained models and minimizes the sum of absolute differences of all enzyme usage values between previous and current time intervals (Materials and Methods). By employing dMORP, we found that the lactate production in P_tre_ was well predicted (Fig. 2B). Moreover, dMORP outperformed other tested objective functions as it led to the smallest Root Mean Square Error (RMSE) (Fig. 2C) and it predicted production of byproducts in P_tre_ such as acetate, acetoin and ethanol (Supplementary Fig. 4). Furthermore, we compared the metabolic fluxes predicted by various objective functions with those by random sampling, which enables unbiased estimations of metabolic fluxes subject to experimentally measured constraints. For reactions in the central carbon metabolism (CCM), the fluxes predicted by dMORP were generally within the ranges of the sampled fluxes, while the fluxes by other objective functions notably deviated (Supplementary Fig. 5). Therefore, we demonstrated that dMORP could be a more realistic objective function for describing the cellular behavior after glucose depletion.

To explain the observed lower lactate yield in P_tre_ with dMORP, we compared the relative fluxes, i.e., normalized to carbon source uptake, of the CCM reactions in P_tre_ (simulated at 48 h, i.e., middle in P_tre_) to those in P_gluc_ (simulated at 7 h, i.e., mid-exponential phase in P_gluc_). We found in P_tre_ a much greater fraction of carbon flux toward phosphoketolase (Fig. 2D), resulting in carbon loss in the form of acetate and ethanol and thereby the decreased lactate yield(14). This was also similar with the comparison between P_tre_ and the sole trehalose condition (Supplementary Fig. 6). In addition, we compared the absolute fluxes simulated at 7 h, 14 h and 48 h, representing P_gluc_, P_lag_ and P_tre_, respectively (Fig. 3). We found that the metabolic fluxes in P_lag_ and P_tre_ were comparable and generally less than those in P_gluc_ (Fig. 3), exemplified by the glycolytic fluxes, which was due to the considerably slower substrate uptake (Supplementary Table 1). By comparing P_gluc_ and P_tre_, we found that the phosphoketolase pathway displayed smaller decreases in fluxes than glycolytic reactions (Supplementary Fig. 7) and thus accounted for a greater fraction of carbon flux, which explained the higher yields of acetate and ethanol.

**Figure 3.**
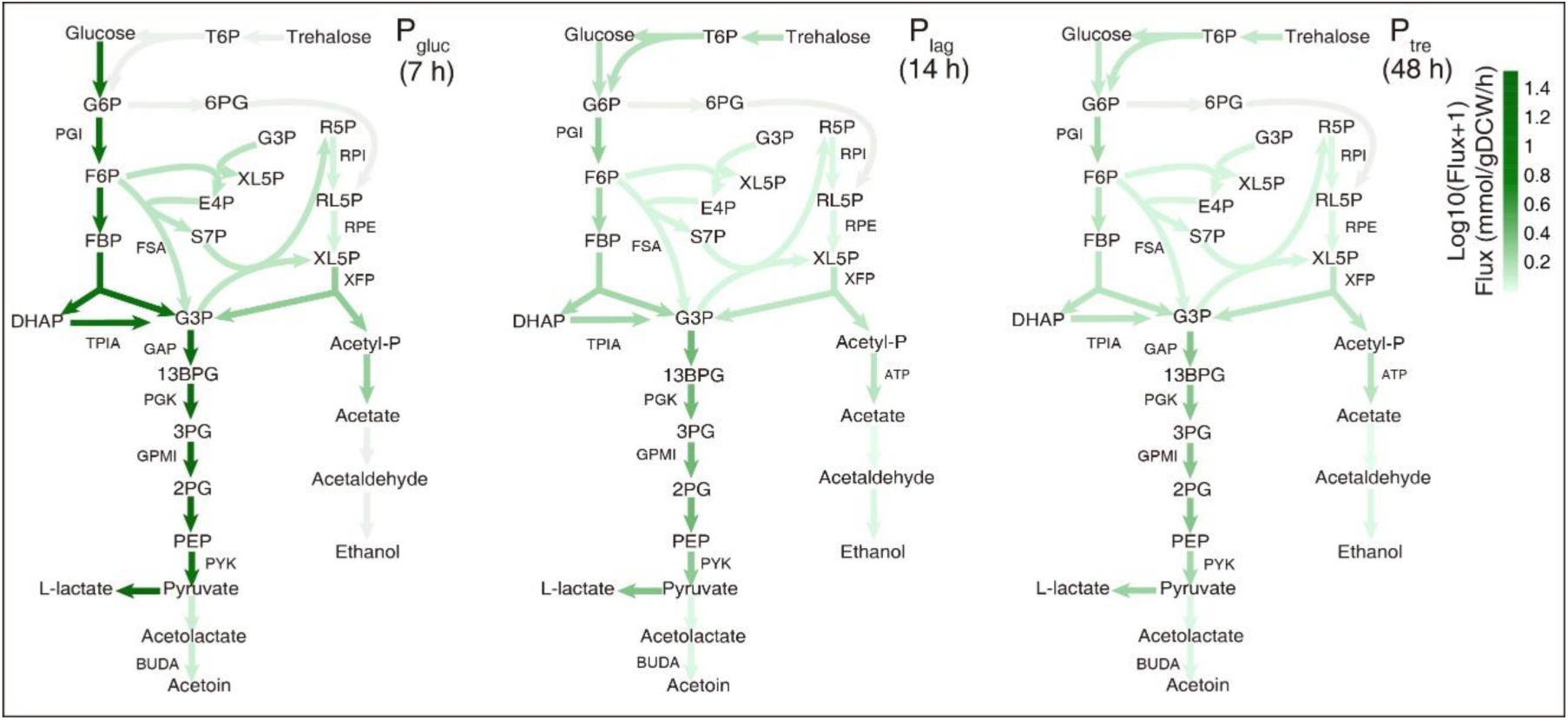
Fluxes of the CCM reactions in the fermentation process predicted by eciBcoa620. The metabolic flux distributions of the CCM at 7 h, 14 h, and 48 h. These three time points represent P_gluc_, P_lag_ and P_tre_, respectively. At 7 h the objective function was to maximize growth while at 14 h and 48 h it was to minimize the proteome reallocation.

### Omics data coincided with dynamic minimization of proteome reallocation

To systematically unravel the molecular mechanisms, we collected transcriptomics and proteomics data at 7 h, 14 h and 48 h, representing P_gluc_, P_lag_ and P_tre_, respectively. The transcriptomics and proteomics data, which showed high consistency among biological triplicates while clear variability among different phases (Supplementary Fig. 8), were used for identifying differentially expressed transcripts and proteins (Materials and Methods). Enrichment analysis of the differential expression showed that from P_gluc_ to P_lag_ and P_tre_ pathways were mostly downregulated, particularly those related to translation, e.g., ribosome and aminoacyl-tRNA biosynthesis (Supplementary Fig. 9), in line with the observed growth arrest (Fig. 1A). Moreover, the differential expression was enriched in amino acid pathways, which could be associated with the significant changes in the concentrations of intracellular amino acids (Supplementary Fig. 10).

Given that the enzyme-constrained models can simulate enzyme usage for metabolic reactions, we compared the simulated enzyme usage to the measured mRNA and protein levels for CCM reactions. We simulated dynamic changes in the enzyme usage using dMORP and dFBA with maximization of growth, lactate and NGAM, and found that dMORP resulted in the least decrease in the enzyme usage upon glucose depletion (Fig. 4), indicating the effectiveness of MORP in minimizing the proteome reallocation during the transition. Notably, the enzyme usage simulated by MORP captured the measured mRNA and protein levels for CCM reactions much better than those simulated by the other objective functions (Fig. 4).

**Figure 4.**
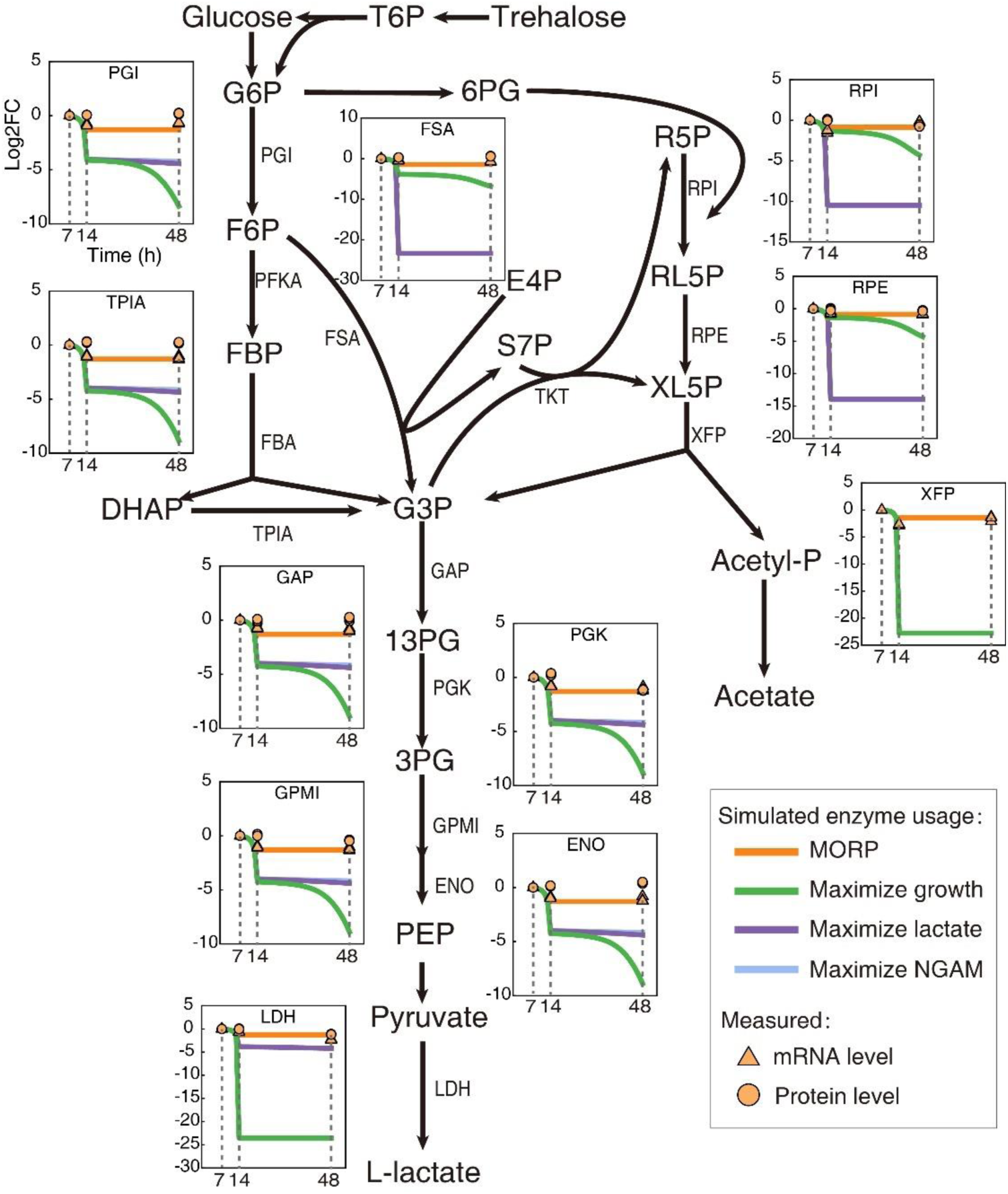
Comparison of simulated enzyme usage and measured mRNA and protein levels. Taking the PGI reaction as an example, the log2FC value represents the change in enzyme usage, mRNA or protein level relative to the 7 h point. The enzyme usage was predicted by different objective functions. The mRNA and protein levels represent average values of biological triplicates from measured transcriptomics and proteomics data.

To further compare with MORP, we performed simulations by constraining exchange rates with measurements and minimizing the total proteome (Materials and Methods), and thus estimated the most efficient usage of the enzymes that maintain the experimentally observed metabolic states. This manner, which is widely adopted for enzyme-constrained simulations(35, 43), also performed worse than MORP in terms of simulating the dynamics of the enzyme usage (Supplementary Fig. 11). In addition, this may indicate suboptimal performance of enzymes during the transition as predicted by MORP (Supplementary Fig. 12), which could be associated with enzyme occupancy by substrates and thermodynamics(44).

In conclusion, the model simulations and experimental measurements demonstrated that the cellular proteome maintained minimal adjustment upon rapid environmental changes, i.e., after glucose depletion cells tended to utilize the proteome expressed on glucose phase to consume trehalose.

### Application of MORP to evolved strains capable of co-utilizing mixed carbon sources

Given that the hierarchical utilization of carbon sources was accompanied by the metabolic transition from homolactic to heterolactic fermentation, we expected that co-utilization of carbon sources would avoid heterolactic fermentation and thus enhance lactate yield. To this end, we carried out ALE (Materials and Methods) (Fig. 5A) and obtained four evolved strains that can co-utilize glucose and trehalose (Supplementary Fig. 13). Whole-genome sequencing identified the mutation in the gene *crr* encoding glucose-specific phosphotransferase enzyme IIA component in all evolved strains (Supplementary Table 2), in line with the findings of the microbes with diminished glucose repression(45–47).

**Figure 5.**
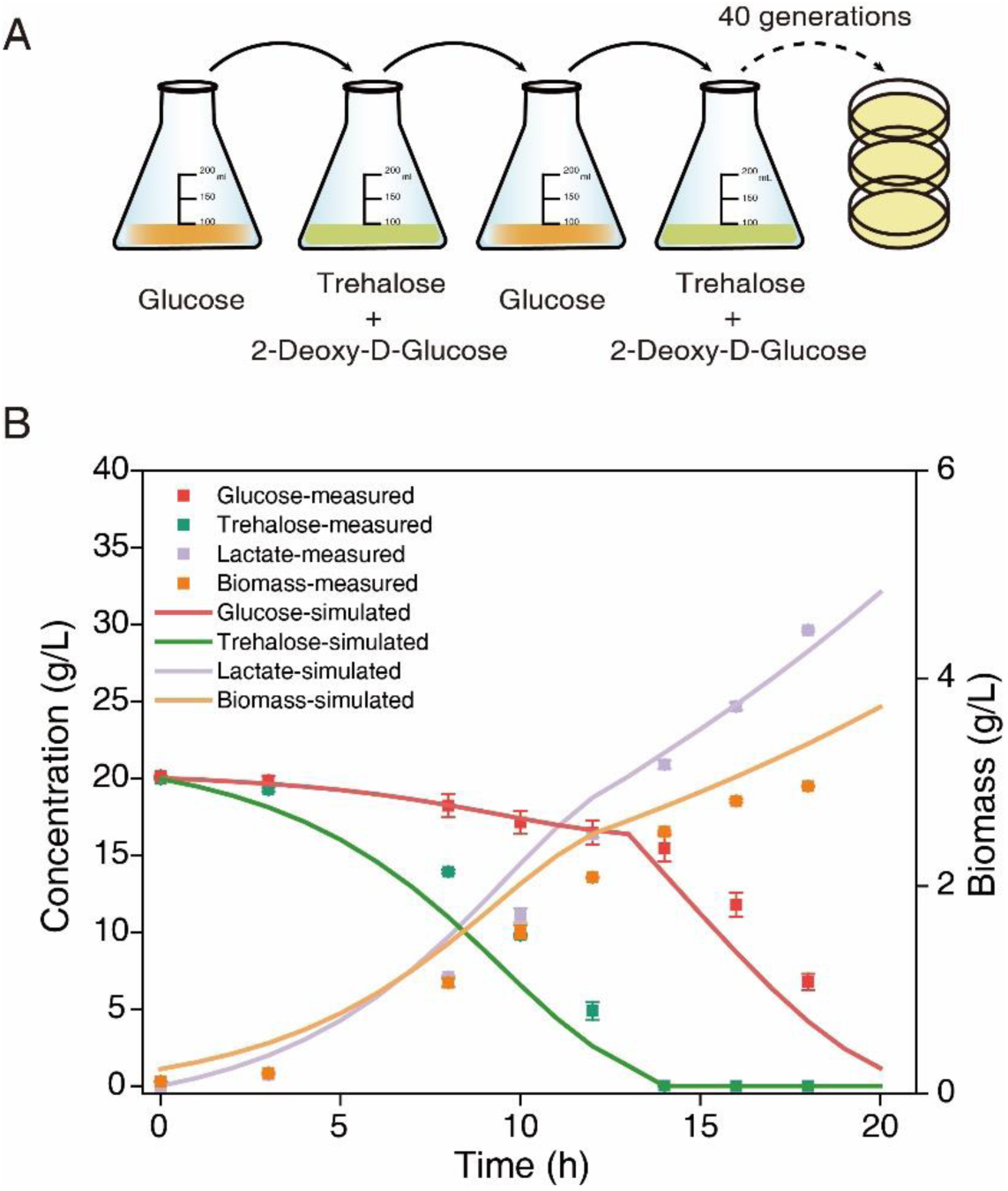
Adaptive laboratory evolution and fermentation process of the evolved strain. (A) Brief diagram of adaptive laboratory evolution experiment. The addition of 2-Deoxy-D-glucose to trehalose aimed at efficiently utilizing trehalose in the absence of glucose, with the intention of eliminating the catabolite repression effect. After 40 generations, growth and lactate production were basically stable. The strains with notably improved lactate production were selected for subsequent shake flasks validation and 5 L bioreactor experiments. (B) Fermentation with mixed carbon sources in 5 L bioreactor by the evolved strain Ev3. Simulations were performed for the fermentation process, in which dFBA with maximization of growth was performed in the phase of simultaneous utilization of glucose and trehalose and dMORP was performed after trehalose depletion. Data are shown with mean ± s.d. of three biological replicates.

We grew one of the evolved strains Ev3 on the mixture of glucose and trehalose in the 5 L bioreactor cultivation and monitored biomass and extracellular metabolite levels (Supplementary Fig. 14). We found that Ev3 utilized glucose and trehalose simultaneously until 14 h when trehalose was exhausted, and then glucose served as the sole carbon source. By calculating the lactate yield for these two phases, we found that Ev3 performed homolactic fermentation on both phases, which led to a higher total lactate yield than the wildtype strain (Supplementary Fig. 14).

Furthermore, the dynamics from mixed carbon sources to glucose utilization provided another scenario that could test MORP. To this end, we simulated the fermentation process of Ev3 within the dFBA framework, in which the first phase (before 14 h) was simulated by maximizing growth and the second (after 14 h) by MORP. We showed that MORP could also capture the physiological changes after trehalose depletion (Fig. 5B), and thus foresee more applications of MORP.

## Discussion

Here, we observed that *B. coagulans* utilized glucose and trehalose hierarchically and transitioned from homolactic to heterolactic fermentation during the hierarchical utilization (Fig. 1). We hypothesized that the dynamic minimization of proteome reallocation might result in this observation. To test it, we reconstructed the enzyme-constrained genome-scale metabolic model of *B. coagulans*, and developed MORP and its dynamic version dMORP. The model with dMORP captured the transition from homolactic to heterolactic fermentation (Fig. 2) and simulated minimal adjustment of proteome during the transition as indicated by the omics data (Fig. 4). We therefore concluded that the dynamic minimization of proteome reallocation could explain the microbial choice of the metabolic strategies upon environmental changes, e.g., changes in the availability of carbon sources in the medium. Last, with the evolved strain we used dMORP to simulate the transition from the co-utilization of glucose and trehalose to the utilization of glucose after trehalose depletion (Fig. 5), demonstrating the extended applications of MORP and dMORP.

The biological basis of the minimization of proteome reallocation is the high cost and slow adjustment of protein synthesis and degradation(42, 48, 49) that the cells should consider when adapting to rapid perturbations. Additionally, the minimization of proteome reallocation is in line with the findings of proteome reserves(50, 51), i.e., the proteome is not fully optimized for the environmental condition where the cells are living, as a suboptimal proteome could reduce the cost and time of reallocation upon the adaption to a new condition. By integrating the minimization of proteome reallocation into the constraint-based modeling framework, we developed MORP and dMORP that introduce the past proteome as an internal constraint, which provide additional objective functions for simulating cellular behaviors(52). Particularly, dMORP would serve as a promising algorithm in the field of dynamic metabolic modeling to predict cellular kinetics(53, 54) and history-dependent behaviors(48, 55, 56).

The approach of minimization of metabolic adjustment (MOMA) has been widely used to simulate cellular behaviors in response to genetic or environmental perturbations(57–59) by minimizing the sum of absolute changes in metabolic fluxes before and after perturbations(60). Compared with minimizing the adjustment of metabolic fluxes, we argue that it is more biologically meaningful to minimize the adjustment of proteome, which now can be achieved by MORP using enzyme-constrained models. It should be noted that MORP can predict the mismatch between enzyme and flux levels, which appears to be an impossible mission for the MOMA approach implemented on conventional GEMs. In addition, it is less computationally expensive to perform the dynamic simulations on enzyme-constrained models than on fine-grained models, such as ME-models(61, 62) that explicitly formulate gene expression processes(26, 63). Therefore, we expect extensive applications of MORP and dMORP with the development of enzyme-constrained models(64).

In summary, we applied systems biology approaches to decipher the principles of the cellular response of *B. coagulans* upon the hierarchical utilization of mixed carbon sources, and identified the dynamic minimization of proteome reallocation to be the cellular objective that dominates the metabolic behavior in response to rapid environmental changes. In addition, we implemented the minimization of proteome reallocation as an effective objective function in the metabolic modeling framework, which would be useful for simulating and understanding cellular responses upon perturbations.

## Materials and Methods

### Microorganism, media and culture conditions

*B. coagulans* DSM 1 = ATCC 7050 was purchased from the American Type Culture Collection. Seed medium was MRS medium(65), simplified chemically defined medium (CDM) used for fermentation in this study, i.e., MCDM3++, which was developed in our previous study(33). 20 g/L glucose and 20 g/L trehalose were used as the mixed carbon source to better reproduce the phenomenon of hierarchical utilization.

Seed culture and shake flask fermentation were performed in 250 mL shake flasks with 100 mL working volume at 50 °C and 100 rpm, pH was buffered with CaCO_3_. The cells in seed culture were collected and washed twice using 100 mM phosphate buffer at around 12 h. Collected cells were resuspended in ultrapure water with a determined volume and then transferred to the fermentation medium in shake flasks or 5 L bioreactors. Shake flask fermentation was used to investigate the metabolic properties of different combinations of mixed carbon sources.

Batch fermentation was performed in a 5 L bioreactor with 4 L working volume at 50 °C, 100 rpm and 7.2 L/h aeration, and pH was automatically controlled at 5.5 using 25% w/v Ca(OH)_2_. The samples of different phases were collected for determination of biomass, metabolites, transcriptome and proteome. Fermentations were performed in triplicates for all conditions.

### Determination of biomass and metabolites

The optical density (OD) was detected using a spectrophotometer at 620 nm to characterize the biomass concentration.

Lactate, acetate, citate, pyruvate and acetoin in the fermentation broth were determined by high performance liquid chromatography (HPLC) (Shimadzu LC-20AT)(13). The Hi-Plex H (300 mm × 7.7 mm) column was used with 210 nm wavelength and 50 °C column temperature; the mobile phase was 0.01 mol/L sulfuric acid solution and the flow rate was set to 0.4 mL/min. The HPLC detection method for ethanol is as follows: the Aminex HPX-87H (300 mm × 7.8 mm) column was used with refractive index detector and 60 °C column temperature; the mobile phase was 0.01 mol/L sulfuric acid solution and the flow rate was set to 0.6 mL/min.

The carbon sources in the fermentation broth were determined by ion chromatography (Dionex ICS-3000) with Dionex Carbopac PA 20 column, the method was referred to the previous description(30) and modified. The pretreatment process of the samples was performed as follows: 1) The samples were diluted to achieve sugar concentrations within the detection range. 2) Na_2_C_2_O_4_ was used to precipitate calcium ions in the culture medium. 3) Proteins were removed from the samples using ethanol. 4) The processed solution was evaporated to obtain the solid form of the sugar, which was then dissolved in water to obtain the sample for testing. The mobile phase was ultrapure water and 200 mmol/L NaOH, respectively. The elution procedure was ultrapure water and 200 mmol/L NaOH eluted at 9:1 ratio for 18 min, then adjusted the ratio to 2:8 to elute for 17 min, and finally adjusted to 9:1 to balance the column for 35 min, and the flow rate for the entire process was set to 0.4 mL/min.

### Reconstruction of genome-scale metabolic models

The model iBcoa620 was reconstructed based on the first *B. coagulans* model iBag597(14): 1) A combined draft model from KEGG(66) and MetaCyc(67) pathway databases based on the genome sequences of *B. coagulans* DSM 1 = ATCC 7050 (GCF_000832905.1) was reconstructed by RAVEN 2.4.0(68). 2) Another draft model was reconstructed by ModelSEED(69) database based on the genome annotation of RAST(70). 3) Shared reactions in the draft models were integrated with iBag597. 4) The IDs of genes, metabolites, and reactions were unified into the form of the KEGG database. 5) Reactions with mass imbalance were corrected. 6) The memote score system was used to assess the model quality. 7) The model was validated by comparing the predicted and measure growth rates under different carbon sources.

The enzyme-constrained model eciBcoa620 was reconstructed using the GECKO toolbox 3.0. Note that *k*_cat_ values were obtained from BRENDA(71) and GotEnzymes(72) databases and predicted using DLKcat tool(73). Molecular weights of all enzymes were obtained from UniProt(74) database.

### Simulation methods

MORP is implemented on the enzyme-constraint models, which can estimate both metabolic and enzyme usage fluxes. MORP estimates the metabolic and enzyme usage fluxes of the model upon a perturbation based on the minimization of proteome reallocation. Thus, the fluxes of the model from a reference condition should be used as the input of MORP, which can be obtained in advance by other algorithms, e.g., FBA, with the enzyme-constraint models. The core of MORP is to minimize the sum of absolute differences of enzyme usage fluxes between the reference and the perturbed conditions:

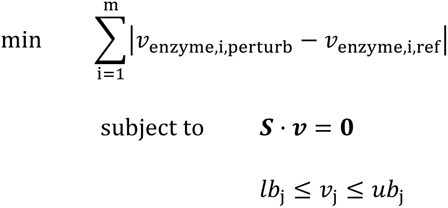

where *v*_enzyme,i,perturb_ and *v*_enzyme,i,ref_ represent fluxes of enzyme usage reaction i under perturbed and reference conditions, and m represents the total number of enzyme usage reactions in the model. As MORP estimates fluxes for the perturbed condition, ***S*** and ***v*** represent the stoichiometric matrix and flux vector of the model for the perturbed condition, and *v*_j_, *lb*_j_ and *ub*_j_ are the flux, lower and upper bounds of the reaction j, respectively.

dFBA based on static optimization approach (75) was used to simulate for the hierarchical utilization of glucose and trehalose, in which an objective function should be determined for different time intervals. As mentioned before three objective functions were used, i.e., maximization of growth, lactate production and NGAM. Generally, dFBA solves:

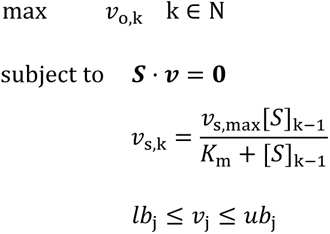

where k represents the k_th_ time interval, N represents the number of time intervals of the phase of interest and thus is a positive integer, *v*_o,k_ is the flux of the objective reaction in P_gluc_, i.e., the biomass formation, or the flux of the objective reaction after glucose depletion, i.e., the biomass formation, lactate production, or NGAM production, *v*_s,k_ is the flux of the uptake reaction of the sugar, i.e., glucose or trehalose, of the kth time interval, [*S*]_k-1_ is the sugar concentration of the (k- 1)_th_ time interval, *v*_s,max_ is the maximum sugar uptake rate, and *K*_m_ is the sugar saturation constant. The constraints in dFBA were derived from experimentally measured sugars, amino acids, organic acids data.

dMORP was proposed to simulate the entire period after glucose depletion in the hierarchical utilization of glucose and trehalose. Here dMORP solves:

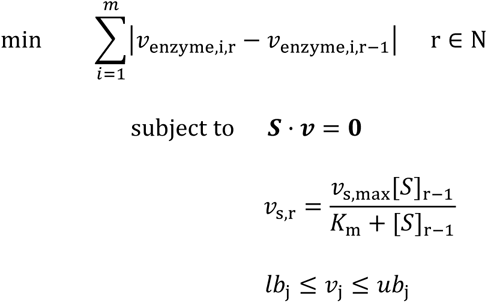

where r represents the r_th_ time interval after glucose depletion, N represents the number of time intervals of the phase of interest and thus is a positive integer, *v*_enzyme,i,r_ and *v*_enzyme,i,r-1_ represent the flux of enzyme usage reaction i for the r_th_ and (r-1)_th_ time interval, respectively, *v*_s,r_ is the flux of the trehalose uptake reaction of the r_th_ time interval, and [*S*]_r-1_ is the trehalose concentration of the (r- 1)_th_ time interval.

Minimization of the total proteome was performed to estimate the most efficient usage of the enzymes that maintain the experimentally observed metabolic states:

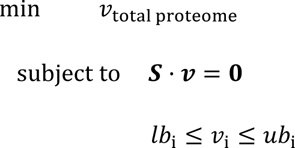

where *v*_total proteome_ represents the flux of total proteome usage, *v*_i_ represents the flux of exchange reaction of detected extracellular metabolite, *lb*_i_ and *ub*_i_ are lower and upper bounds based on measurements.

All the simulations were performed with the COBRA toolbox 3.0(76) on Matlab R2019a, and the solver was Gurobi 9.1 (https://www.gurobi.com/).

### Transcriptome analysis

We collected transcriptomics samples at 7 h, 14 h and 48 h, representing P_gluc_, P_lag_ and P_tre_, respectively. Cells in the fermentation broth were collected and washed three times with phosphate buffer saline for subsequent RNA extraction. Total RNA was isolated from cells using TRIzol Reagent following the instructions and genomic DNA was removed using DNase I (TaKara). Subsequently, RNA quality was determined using a Agilent2100 Bioanalyzer and quantified using ND-2000 (NanoDrop Technologies).

RNA library construction was performed using TruSeqTM RNA sample preparation Kit from Illumina (San Diego, CA). The rRNA was removed using Ribo-Zero Magnetic kit (epicenter), and double-stranded cDNA was reverse transcribed using random primers (Illumina) and SuperScript double-stranded cDNA synthesis kit (Invitrogen, CA). To synthesize the second strand of cDNA, it uses dUTP instead of dTTP, and then eliminates the second strand of cDNA containing dUTP, so that the library contained only the first strand of cDNA. After PCR amplification by Phusion DNA polymerase (NEB) and TBS380 (Picogreen) quantification, paired-end RNA-seq sequencing library was sequenced with the Illumina Novaseq (2 × 150bp read length).

The data generated from Illumina platform were used for bioinformatics analysis, and the reference genome and gene annotation files of *B. coagulans* DSM 1 = ATCC 7050 were downloaded from the National Center for Biotechnology Information. All of the analyses were performed using the free online platform of Majorbio Cloud Platform (www.majorbio.com).

### Proteome analysis

We collected proteomics samples at 7 h, 14 h and 48 h, representing P_gluc_, P_lag_ and P_tre_, respectively. Cells in the fermentation broth were collected and washed three times with phosphate buffer saline for subsequent protein extraction. Sample lysis and protein extraction were performed using SDT (4% SDS, 100 mM Tris-HCl, 1 mM DTT, pH 7.6) buffer, and the protein was quantified using the BCA Protein Assay Kit (Bio-Rad, USA). Protein digestion was performed with trypsin following the filter-aided sample preparation (FASP) procedure. The digested peptides from each sample were subsequently desalted (Empore™ SPE Cartridges C18 (standard density), bed I.D. 7 mm, volume 3 mL, Sigma) and concentrated. The protein for each sample were quantified by SDS-PAGE. 100 μg peptide mixture for each sample was labeled using TMT reagent according to the instructions (Thermo Scientific). The labeled peptides were separated by High pH Reversed-Phase Peptide Fractionation Kit (Thermo Scientific) to obtain 10 different fractions, the collected fractions were desalted on C18 Cartridges and concentrated.

LC-MS/MS analysis was performed using a Thermo Scientific Q Exactive mass spectrometer equipped with the Easy nLC system (Proxeon Biosystems, Thermo Fisher Scientific). Peptides were trapped on a reverse phase trap column (Thermo Scientific Acclaim PepMap100, 100 μm × 2 cm, nanoViper C18) and separated on the C18-reversed phase analytical column (Thermo Scientific Easy Column, 10 cm long, 75 μm inner diameter, 3 μm resin). The separation was achieved using buffer A (0.1% formic acid) and buffer B (84% acetonitrile and 0.1% formic acid) at a constant flow rate of 300 nL/min controlled by IntelliFlow technology. The mass spectrometer was operated in positive ion mode. MS/MS analysis was performed using the data-dependent top10 method for high-energy collision dissociation (HCD) fragmentation by selecting the most abundant precursor ion (300-1800 m/z) in the measurement scan. The MS raw data was subjected to identification and quantitative analysis using MASCOT engine (Matrix Science, London, UK; version 2.2) and Proteome Discoverer 1.4 software. The proteome detection and analysis process were conducted in shanghai applied protein technology co.ltd.

### Differential expression analysis

Significance analysis between two conditions or phases were performed using the two-sided *t*-test, and a cut-off adjusted *P* value (*P*_adjusted_) of 0.01 was adopted for identifying differentially expressed transcripts and proteins based on transcriptomics and proteomics data.

### Determination of intracellular metabolites

The absolute quantification of intracellular amino acids was performed using gas chromatography-mass spectrometry (Agilent, GC-7890A, MS-5975C). Before the extraction of intracellular metabolites, a quenching step was performed to terminate cell metabolism. 2 mL of fermentation broth were added to a 50 mL centrifuge tube containing 30 mL of 60% methanol pre-cooled to - 27 °C. After quenching for 5 minutes, filter paper with attached cells was obtained by filtration. Simultaneously, 100 μL of U-^13^C-labeled cell extracts was added to the filter paper as internal standards. The filter paper was then placed in a 75% ethanol solution pre-heated to 80 °C and boiled for 5 min. The filter paper was removed, and the remaining liquid was concentrated to 600 μL using rapid evaporation. A 100 μL portion was taken as the sample and subjected to freeze-drying before derivatization(14, 77, 78).

Amino acids derivatization method: 100 μL of pyridine was added to the freeze-dried sample, and the mixture was dissolved in a 65 °C for 1 hour. After the sample reached room temperature, 100 μL of derivatization reagent (N - tert-butyldimethylsilyl - N-methyl-trifluoroacetamide with 1% tert - butyldimethylchlorosilane) was added, and derivatization was carried out in a 65 °C oven for 1 hour. The sample was then centrifuged at 12000 rpm for 5 min, and the supernatant was subjected to GC-MS analysis.

The GC-MS detection conditions were as follows: the interface temperature of the GC-MS was set at 280 °C, the transfer line temperature was 250 °C, Ionization was done by electron impact with the temperature being 230 °C, the quadrupole temperature was 150 °C, and the injection volume was 1 μL. The chromatographic column used was HP-5MS with dimensions of 30 mm × 0.25 mm × 0.25 μm. For amino acids, the temperature program consisted of an initial column temperature of 100 °C for 1 minute, followed by an increase at a rate of 10 °C/min to reach 320 °C, which was then maintained for 10 minutes.

### Adaptive laboratory evolution

The adaptive laboratory evolution strategy was performed in a 250 mL shake flask with complex medium, 100 rpm, 50 °C. Alternating cultures of glucose and trehalose were utilized, while the trehalose was supplemented with glucose structural analogue (2-deoxy-D-glucose), which can be absorbed but not be metabolized. Therefore, other sugars are still able to be metabolized even in the presence of glucose, so 2-deoxy-D-glucose can be used for screening mutant strains without catabolite repression effect (79, 80). The aim was to obtain strains that could utilize glucose and trehalose simultaneously. During the evolution process, the cells were transferred to new medium in the exponential phase and maintained the same concentration (OD_620_ about 0.5) after inoculation. The initial total sugar concentration during evolution was kept constant at 130 g/L (glucose 130 g/L, 2-deoxy-D-glucose 40 g/L + trehalose 90 g/L). 40 g/L of 2-deoxy-D-glucose was selected as no growth occurred at higher concentrations. Four evolved strains with the expected phenotypes were examined by MRS plate culture, and the strain with faster sugar consumption rate and higher lactate yield was selected for fermentation experiments in 5 L bioreactor.

### Whole-genome sequencing

Cells from the exponential phase in seed shake flasks were collected for genome sequencing. Genomic DNA of *B. coagulans* DSM 1 = ATCC 7050 was extracted using Wizard Genomic DNA Purification Kit (Promega) according to the protocol. Purified genomic DNA was quantified by TBS- 380 fluorometer (Turner BioSystems Inc., Sunnyvale, CA). Whole-genome sequencing of wildtype strain and four evolved strains was performed using the de novo sequencing method based on the Illumina NovaSeq6000 platform in this study. The final genome of five strains were annotated from KEGG, GO, COG and other databases using BLASTP, Diamond and HMMER. The same mutated genes of four evolved strains were considered to be genes that could cause phenotypic change.

## Data availability

The genome sequences of four evolved strains have been deposited in NCBI database under the BioProject accession number of PRJNA1012373. The transcriptomics data have been deposited in NCBI database under the BioProject accession number of PRJNA1012495. The mass spectrometry proteomics datasets have been deposited in the ProteomeXchange with the dataset identifier PXD045169.

## Code availability

The models and code are available at https://github.com/ChenYuGroup/MORP.

## Acknowledgments

This study was financially supported by the National Key R&D Program of China (2023YFA0913900), the Shenzhen Medical Research Fund (A2303026) and the National Natural Science Foundation of China (21878084).

## Author contributions

Y.C. and Y.W. conceived the study. Z.L., M.C. and J.H. performed the experiments. Z.L. and Y.C. constructed the models and performed the simulations. Z.L. and Y.C. analyzed the data. Y.C., Z.L. and Y.W. wrote the paper. All authors approved the final paper.

## Competing Interest

The authors declare no competing interests.

## Notes

### Competing Interest Statement

The authors have declared no competing interest.

### Summary of Updates

Detailed changes and descriptions were made to the 'omics data coincided with dynamic minimization of proteome reallocation' section，the relevant images have been adjusted.

## References

1. Heiden MGV, Cantley LC, Thompson CB. 2009. Understanding the Warburg Effect: The Metabolic Requirements of Cell Proliferation. Science 324:1029–1033.

2. Shen Y, Dinh HV, Cruz ER, Chen Z, Bartman CR, Xiao T, Call CM, Ryseck R-P, Pratas J, Weilandt D, Baron H, Subramanian A, Fatma Z, Wu Z-Y, Dwaraknath S, Hendry JI, Tran VG, Yang L, Yoshikuni Y, Zhao H, Maranas CD, Wuehr M, Rabinowitz JD. 2024. Mitochondrial ATP generation is more proteome efficient than glycolysis. Nature Chemical Biology doi:10.1038/s41589-024-01571-y.

3. Pfeiffer T, Schuster S, Bonhoeffer S. 2001. Cooperation and competition in the evolution of ATP-producing pathways. Science 292:504–507.

4. Molenaar D, van Berlo R, de Ridder D, Teusink B. 2009. Shifts in growth strategies reflect tradeoffs in cellular economics. Molecular Systems Biology 5:323.

5. Flamholz A, Noor E, Bar-Even A, Liebermeister W, Milo R. 2013. Glycolytic strategy as a tradeoff between energy yield and protein cost. Proceedings of the National Academy of Sciences of the United States of America 110:10039–10044.

6. Szenk M, Dill KA, de Graff AMR. 2017. Why Do Fast-Growing Bacteria Enter Overflow Metabolism? Testing the Membrane Real Estate Hypothesis. Cell Systems 5:95–104.

7. Du B, Zielinski DC, Monk JM, Palsson BO. 2018. Thermodynamic favorability and pathway yield as evolutionary tradeoffs in biosynthetic pathway choice. Proceedings of the National Academy of Sciences of the United States of America 115:11339–11344.

8. Mori M, Marinari E, De Martino A. 2019. A yield-cost tradeoff governs Escherichia coli’s decision between fermentation and respiration in carbon-limited growth. NPJ Syst Biol Appl 5:16.

9. Basan M, Hui S, Okano H, Zhang Z, Shen Y, Williamson JR, Hwa T. 2015. Overflow metabolism in Escherichia coli results from efficient proteome allocation. Nature 528:99–104.

10. Chen Y, Nielsen J. 2019. Energy metabolism controls phenotypes by protein efficiency and allocation. Proceedings of the National Academy of Sciences of the United States of America 116:17592–17597.

11. Schuetz R, Zamboni N, Zampieri M, Heinemann M, Sauer U. 2012. Multidimensional Optimality of Microbial Metabolism. Science 336:601–604.

12. Mori M, Schink S, Erickson DW, Gerland U, Hwa T. 2017. Quantifying the benefit of a proteome reserve in fluctuating environments. Nature Communications 8.

13. Lv X, Yu B, Tian X, Chen Y, Wang Z, Zhuang Y, Wang Y. 2016. Effect of pH, glucoamylase, pullulanase and invertase addition on the degradation of residual sugar in L-lactic acid fermentation by Bacillus coagulans HL-5 with corn flour hydrolysate. Journal of the Taiwan Institute of Chemical Engineers 61:124–131.

14. Chen Y, Sun Y, Liu Z, Dong F, Li Y, Wang Y. 2020. Genome-scale modeling for Bacillus coagulans to understand the metabolic characteristics. Biotechnology and Bioengineering 117:3545–3558.

15. Gaenzle MG. 2015. Lactic metabolism revisited: metabolism of lactic acid bacteria in food fermentations and food spoilage. Current Opinion in Food Science 2:106–117.

16. Wang Y, Wu J, Lv M, Shao Z, Hungwe M, Wang J, Bai X, Xie J, Wang Y, Geng W. 2021. Metabolism Characteristics of Lactic Acid Bacteria and the Expanding Applications in Food Industry. Frontiers in Bioengineering and Biotechnology 9:612285.

17. Wang Q, Ingram LO, Shanmugam KT. 2011. Evolution of D-lactate dehydrogenase activity from glycerol dehydrogenase and its utility for D-lactate production from lignocellulose. Proceedings of the National Academy of Sciences of the United States of America 108:18920–18925.

18. Wang L, Xue Z, Zhao B, Yu B, Xu P, Ma Y. 2013. Jerusalem artichoke powder: A useful material in producing high-optical-purity L-lactate using an efficient sugar-utilizing thermophilic Bacillus coagulans strain. Bioresource Technology 130:174–180.

19. Abdel-Rahman MA, Tashiro Y, Sonomoto K. 2013. Recent advances in lactic acid production by microbial fermentation processes. Biotechnology Advances 31:877–902.

20. Lu Z, He F, Shi Y, Lu M, Yu L. 2010. Fermentative production of L(+)-lactic acid using hydrolyzed acorn starch, persimmon juice and wheat bran hydrolysate as nutrients. Bioresource Technology 101:3642–3648.

21. Hermsen R, Okano H, You C, Werner N, Hwa T. 2015. A growth-rate composition formula for the growth of E. coli on co-utilized carbon substrates. Molecular Systems Biology 11:801.

22. Perrin E, Ghini V, Giovannini M, Di Patti F, Cardazzo B, Carraro L, Fagorzi C, Turano P, Fani R, Fondi M. 2020. Diauxie and co-utilization of carbon sources can coexist during bacterial growth in nutritionally complex environments. Nature Communications 11:3135.

23. Okano H, Hermsen R, Hwa T. 2021. Hierarchical and simultaneous utilization of carbon substrates: mechanistic insights, physiological roles, and ecological consequences. Current Opinion in Microbiology 63:172–178.

24. Solopova A, van Gestel J, Weissing FJ, Bachmann H, Teusink B, Kok J, Kuipers OP. 2014. Bet-hedging during bacterial diauxic shift. Proceedings of the National Academy of Sciences of the United States of America 111:7427–7432.

25. Okano H, Hermsen R, Kochanowski K, Hwa T. 2020. Regulation underlying hierarchical and simultaneous utilization of carbon substrates by flux sensors in Escherichia coli. Nature Microbiology 5:206–215.

26. Salvy P, Hatzimanikatis V. 2021. Emergence of diauxie as an optimal growth strategy under resource allocation constraints in cellular metabolism. Proceedings of the National Academy of Sciences of the United States of America 118:e2013836118.

27. Sievert C, Nieves LM, Panyon LA, Loeffler T, Morris C, Cartwright RA, Wang X. 2017. Experimental evolution reveals an effective avenue to release catabolite repression via mutations in XylR. Proceedings of the National Academy of Sciences of the United States of America 114:7349–7354.

28. Gawand P, Hyland P, Ekins A, Martin VJJ, Mahadevan R. 2013. Novel approach to engineer strains for simultaneous sugar utilization. Metabolic Engineering 20:63–72.

29. Ma K, Hu G, Pan L, Wang Z, Zhou Y, Wang Y, Ruan Z, He M. 2016. Highly efficient production of optically pure L-lactic acid from corn stover hydrolysate by thermophilic Bacillus coagulans. Bioresource Technology 219:114–122.

30. Lv X, Guo Y, Zhuang Y, Wang Y. 2015. Optimization and validation of an extraction method and HPAEC-PAD for determination of residual sugar composition in L- lactic acid industrial fermentation broth with a high salt content. Analytical Methods 7:9076–9083.

31. Wee YJ, Yun JS, Park DH, Ryu HW. 2004. Isolation and characterization of a novel lactic acid bacterium for the production of lactic acid. Biotechnology and Bioprocess Engineering 9:303–308.

32. Dooley D, Ryu S, J. Giannone R, Edwards J, S. Dien B, J. Slininger P, T. Trinh C. 2024. Expanded genome and proteome reallocation in a novel, robust Bacillus coagulans strain capable of utilizing pentose and hexose sugars. mSystems:e00952–24.

33. Chen Y, Dong F, Wang Y. 2016. Systematic development and optimization of chemically defined medium supporting high cell density growth of Bacillus coagulans. Applied Microbiology and Biotechnology 100:8121–8134.

34. Lieven C, Beber ME, Olivier BG, Bergmann FT, Ataman M, Babaei P, Bartell JA, Blank LM, Chauhan S, Correia K, Diener C, Draeger A, Ebert BE, Edirisinghe JN, Faria JP, Feist AM, Fengos G, Fleming RMT, Garcia-Jimenez B, Hatzimanikatis V, van Helvoirt W, Henry CS, Hermjakob H, Herrgard MJ, Kaafarani A, Kim HU, King Z, Klamt S, Klipp E, Koehorst JJ, Koenig M, Lakshmanan M, Lee D-Y, Lee SY, Lee S, Lewis NE, Liu F, Ma H, Machado D, Mahadevan R, Maia P, Mardinoglu A, Medlock GL, Monk JM, Nielsen J, Nielsen LK, Nogales J, Nookaew I, Palsson BO, Papin JA, et al. 2020. MEMOTE for standardized genome-scale metabolic model testing. Nature Biotechnology 38:272–276.

35. Chen Y, Gustafsson J, Tafur Rangel A, Anton M, Domenzain I, Kittikunapong C, Li F, Yuan L, Nielsen J, Kerkhoven EJ. 2024. Reconstruction, simulation and analysis of enzyme-constrained metabolic models using GECKO Toolbox 3.0. Nature Protocols 19:629–667.

36. Alter TB, Blank LM, Ebert BE. 2021. Proteome Regulation Patterns Determine Escherichia coli Wild-Type and Mutant Phenotypes. Msystems 6.

37. Mao Z, Niu J, Zhao J, Huang Y, Wu K, Yun L, Guan J, Yuan Q, Liao X, Wang Z, Ma H. 2024. ECMpy 2.0: A Python package for automated construction and analysis of enzyme-constrained models. Synthetic and systems biotechnology 9:494–502.

38. Ye C, Luo Q, Guo L, Gao C, Xu N, Zhang L, Liu L, Chen X. 2020. Improving lysine production through construction of an Escherichia coli enzyme-constrained model. Biotechnology and Bioengineering 117:3533–3544.

39. Orth JD, Thiele I, Palsson BO. 2010. What is flux balance analysis? Nature Biotechnology 28:245–248.

40. Henriques D, Minebois R, Mendoza SN, Macías LG, Pérez-Torrado R, Barrio E, Teusink B, Querol A, Balsa-Canto E. 2021. A multiphase multiobjective dynamic genome-scale model shows different redox balancing among yeast species of the Saccharomyces genus in fermentation. msystems 6:10–1128.

41. Moreno-Paz S, Schmitz J, Martins Dos Santos VAP, Suarez-Diez M. 2022. Enzyme-constrained models predict the dynamics of Saccharomyces cerevisiae growth in continuous, batch and fed-batch bioreactors. Microb Biotechnol 15:1434–1445.

42. Kafri M, Metzl-Raz E, Jona G, Barkai N. 2016. The Cost of Protein Production. Cell Reports 14:22–31.

43. Domenzain I, Sanchez B, Anton M, Kerkhoven EJ, Millan-Oropeza A, Henry C, Siewers V, Morrissey JP, Sonnenschein N, Nielsen J. 2022. Reconstruction of a catalogue of genome-scale metabolic models with enzymatic constraints using GECKO 2.0. Nature Communications 13.

44. Park JO, Tanner LB, Wei MH, Khana DB, Jacobson TB, Zhang Z, Rubin SA, Li SH-J, Higgins MB, Stevenson DM, Amador-Noguez D, Rabinowitz JD. 2019. Near-equilibrium glycolysis supports metabolic homeostasis and energy yield. Nature Chemical Biology 15:1001–1008.

45. Oh B-R, Hong W-K, Heo S-Y, Luo LH, Kondo A, Seo J-W, Kim CH. 2013. The production of 1,3-propanediol from mixtures of glycerol and glucose by a Klebsiella pneumoniae mutant deficient in carbon catabolite repression. Bioresource Technology 130:719–724.

46. Zhao M, Shi D, Lu X, Zong H, Zhuge B. 2019. Co-production of 1,2,4-butantriol and ethanol from lignocellulose hydrolysates. Bioresource Technology 282:433–438.

47. Liang Q, Zhang F, Li Y, Zhang X, Li J, Yang P, Qi Q. 2015. Comparison of individual component deletions in a glucose-specific phosphotransferase system revealed their different applications. Scientific Reports 5:13200.

48. Rothman DL, Moore PB, Shulman RG. 2023. The impact of metabolism on the adaptation of organisms to environmental change. Frontiers in Cell and Developmental Biology 11:1197226.

49. Scott M, Hwa T. 2023. Shaping bacterial gene expression by physiological and proteome allocation constraints. Nature Reviews Microbiology 21:327–342.

50. Wu C, Mori M, Abele M, Banaei-Esfahani A, Zhang Z, Okano H, Aebersold R, Ludwig C, Hwa T. 2023. Enzyme expression kinetics by Escherichia coli during transition from rich to minimal media depends on proteome reserves. Nature Microbiology 8:347–359.

51. Yu R, Campbell K, Pereira R, Bjorkeroth J, Qi Q, Vorontsov E, Sihlbom C, Nielsen J. 2020. Nitrogen limitation reveals large reserves in metabolic and translational capacities of yeast. Nature Communications 11:1881.

52. Bernstein DB, Sulheim S, Almaas E, Segre D. 2021. Addressing uncertainty in genome-scale metabolic model reconstruction and analysis. Genome Biology 22:1–22.

53. Erickson DW, Schink SJ, Patsalo V, Williamson JR, Gerland U, Hwa T. 2017. A global resource allocation strategy governs growth transition kinetics of Escherichia coli. Nature 551:119–123.

54. Basan M, Honda T, Christodoulou D, Horl M, Chang Y-F, Leoncini E, Mukherjee A, Okano H, Taylor BR, Silverman JM, Sanchez C, Williamson JR, Paulsson J, Hwa T, Sauer U. 2020. A universal trade-off between growth and lag in fluctuating environments. Nature 584:470–474.

55. Cerulus B, Jariani A, Perez-Samper G, Vermeersch L, Pietsch JMJ, Crane MM, New AM, Gallone B, Roncoroni M, Dzialo MC, Govers SK, Hendrickx JO, Galle E, Coomans M, Berden P, Verbandt S, Swain PS, Verstrepen KJ. 2018. Transition between fermentation and respiration determines history-dependent behavior in fluctuating carbon sources. Elife 7.

56. Vermeersch L, Cool L, Gorkovskiy A, Voordeckers K, Wenseleers T, Verstrepen KJ. 2022. Do microbes have a memory? History-dependent behavior in the adaptation to variable environments. Frontiers in Microbiology 13:1004488.

57. Yu JSL, Correia-Melo C, Zorrilla F, Herrera-Dominguez L, Wu MY, Hartl J, Campbell K, Blasche S, Kreidl M, Egger A-S, Messner CB, Demichev V, Freiwald A, Mulleder M, Howell M, Berman J, Patil KR, Alam MT, Ralser M. 2022. Microbial communities form rich extracellular metabolomes that foster metabolic interactions and promote drug tolerance. Nature Microbiology 7:542–555.

58. Ho W-C, Zhang J. 2018. Evolutionary adaptations to new environments generally reverse plastic phenotypic changes. Nature Communications 9:350.

59. Vaud S, Pearcy N, Hanzevacki M, Van Hagen AMW, Abdelrazig S, Safo L, Ehsaan M, Jonczyk M, Millat T, Craig S, Spence E, Fothergill J, Bommareddy RR, Colin P-Y, Twycross J, Dalby PA, Minton NP, Jaeger CM, Kim D-H, Yu J, Maness P-C, Lynch S, Eckert CA, Conradie A, Bryan SJ. 2021. Engineering improved ethylene production: Leveraging systems biology and adaptive laboratory evolution. Metabolic Engineering 67:308–320.

60. Segrè D, Vitkup D, Church GM. 2002. Analysis of optimality in natural and perturbed metabolic networks. Proceedings of the National Academy of Sciences of the United States of America 99:15112–15117.

61. Lloyd CJ, Ebrahim A, Yang L, King ZA, Catoiu E, O’Brien EJ, Liu JK, Polsson BO. 2018. COBRAme: A computational framework for genome-scale models of metabolism and gene expression. Plos Computational Biology 14:e1006302.

62. Reimers A-M, Knoop H, Bockmayr A, Steuer R. 2017. Cellular trade-offs and optimal resource allocation during cyanobacterial diurnal growth. Proceedings of the National Academy of Sciences of the United States of America 114:E6457–E6465.

63. Yang L, Ebrahim A, Lloyd CJ, Saunders MA, Palsson BO. 2019. DynamicME: dynamic simulation and refinement of integrated models of metabolism and protein expression. Bmc Systems Biology 13:1–15.

64. Chen Y, Nielsen J. 2021. Mathematical modeling of proteome constraints within metabolism. Current Opinion in Systems Biology 25:50–56.

65. David AN, Sewsynker-Sukai Y, Kana EBG. 2022. Co-valorization of corn cobs and dairy wastewater for simultaneous saccharification and lactic acid production: Process optimization and kinetic assessment. Bioresource Technology 348.

66. Kanehisa M, Goto S. 2000. KEGG: Kyoto Encyclopedia of Genes and Genomes. Nucleic Acids Research 28:27–30.

67. Karp PD, Riley M, Paley SM, Pellegrini-Toole A. 2002. The MetaCyc database. Nucleic Acids Research 30:59–61.

68. Wang H, Marcisauskas S, Sanchez BJ, Domenzain I, Hermansson D, Agren R, Nielsen J, Kerkhoven EJ. 2018. RAVEN 2.0: A versatile toolbox for metabolic network reconstruction and a case study on Streptomyces coelicolor. Plos Computational Biology 14:e1006541.

69. Henry CS, DeJongh M, Best AA, Frybarger PM, Linsay B, Stevens RL. 2010. High-throughput generation, optimization and analysis of genome-scale metabolic models. Nature Biotechnology 28:977–U22.

70. Aziz RK, Bartels D, Best AA, DeJongh M, Disz T, Edwards RA, Formsma K, Gerdes S, Glass EM, Kubal M, Meyer F, Olsen GJ, Olson R, Osterman AL, Overbeek RA, McNeil LK, Paarmann D, Paczian T, Parrello B, Pusch GD, Reich C, Stevens R, Vassieva O, Vonstein V, Wilke A, Zagnitko O. 2008. The RAST server: Rapid annotations using subsystems technology. Bmc Genomics 9.

71. Jeske L, Placzek S, Schomburg I, Chang A, Schomburg D. 2019. BRENDA in 2019: a European ELIXIR core data resource. Nucleic Acids Research 47:D542–D549.

72. Li F, Chen Y, Anton M, Nielsen J. 2023. GotEnzymes: an extensive database of enzyme parameter predictions. Nucleic Acids Research 51:D583–D586.

73. Li F, Yuan L, Lu H, Li G, Chen Y, Engqvist MKM, Kerkhoven EJ, Nielsen J. 2022. Deep learning-based kcat prediction enables improved enzyme-constrained model reconstruction. Nature Catalysis 5:662–672.

74. Apweiler R, Bairoch A, Wu CH, Barker WC, Boeckmann B, Ferro S, Gasteiger E, Huang HZ, Lopez R, Magrane M, Martin MJ, Natale DA, O’Donovan C, Redaschi N, Yeh LSL. 2004. UniProt: the Universal Protein knowledgebase. Nucleic Acids Research 32:D115–D119.

75. Mahadevan R, Edwards JS, Doyle FJ. 2002. Dynamic flux balance analysis of diauxic growth in Escherichia coli. Biophysical Journal 83:1331–1340.

76. Heirendt L, Arreckx S, Pfau T, Mendoza SN, Richelle A, Heinken A, Haraldsdottir HS, Wachowiak J, Keating SM, Vlasov V, Magnusdottir S, Ng CY, Preciat G, Zagare A, Chan SHJ, Aurich MK, Clancy CM, Modamio J, Sauls JT, Noronha A, Bordbar A, Cousins B, El Assal DC, Valcarcel LV, Apaolaza I, Ghaderi S, Ahookhosh M, Ben Guebila M, Kostromins A, Sompairac N, Le HM, Ma D, Sun Y, Wang L, Yurkovich JT, Oliveira MAP, Vuong PT, El Assal LP, Kuperstein I, Zinovyev A, Hinton HS, Bryant WA, Aragon Artacho FJ, Planes FJ, Stalidzans E, Maass A, Vempala S, Hucka M, Saunders MA, Maranas CD, et al. 2019. Creation and analysis of biochemical constraint-based models using the COBRA Toolbox v.3.0. Nature Protocols 14:639–702.

77. Wang G, Chu J, Zhuang Y, van Gulik W, Noorman H. 2019. A dynamic model-based preparation of uniformly-13C-labeled internal standards facilitates quantitative metabolomics analysis of Penicillium chrysogenum. Journal of Biotechnology 299:21–31.

78. Liu X, Sun X, He W, Tian X, Zhuang Y, Chu J. 2019. Dynamic changes of metabolomics and expression of candicidin biosynthesis gene cluster caused by the presence of a pleiotropic regulator AdpA in Streptomyces ZYJ-6. Bioprocess and Biosystems Engineering 42:1353–1365.

79. Liang J, van Kranenburg R, Bolhuis A, Leak DJ. 2022. Removing carbon catabolite repression in Parageobacillus thermoglucosidasius DSM 2542. Frontiers in Microbiology 13:985465.

80. Veyrat A, Monedero V, Perez-Martinez G. 1994. Glucose transport by the phosphoenolpyruvate: mannose phosphotransferase system in Lactobacillus casei ATCC 393 and its role in carbon catabolite repression. Microbiology 140:1141–1149.

