## supplemental information for "Minimization of proteome reallocation explains metabolic transition in hierarchical utilization of carbon sources"

### This file includes:

### Supplementary Fig. 1

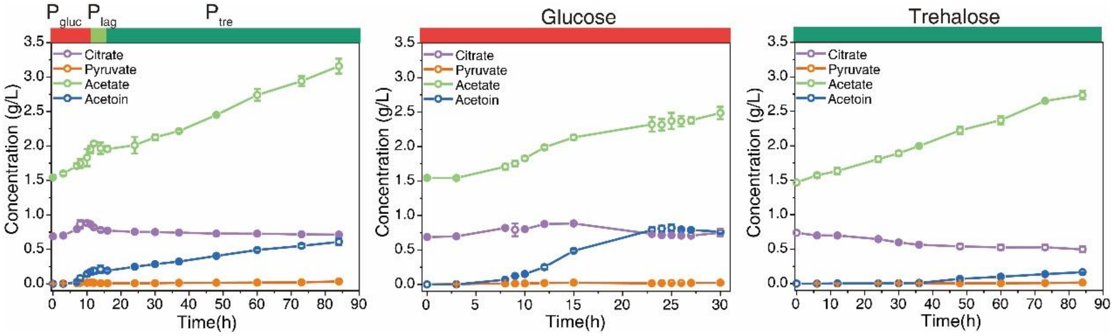

**Supplementary Fig. 1. Fermentation phenotypes under different carbon source conditions in 5 L bioreactor.**

Process profiles of byproduct organic acids under different carbon source conditions. Under the mixed carbon source condition, organic acids production followed a two-phase pattern consistent with hierarchical utilization, some organic acids being utilized during the transition between these phases to cope with the carbon sources gap. Both sole glucose and sole trehalose exhibited smooth curves in organic acids production, in sharp contrast to the distinct pattern observed under the mixed carbon source. Data in point line diagram are mean ± s.d. of three biological replicates.

### Supplementary Fig. 2

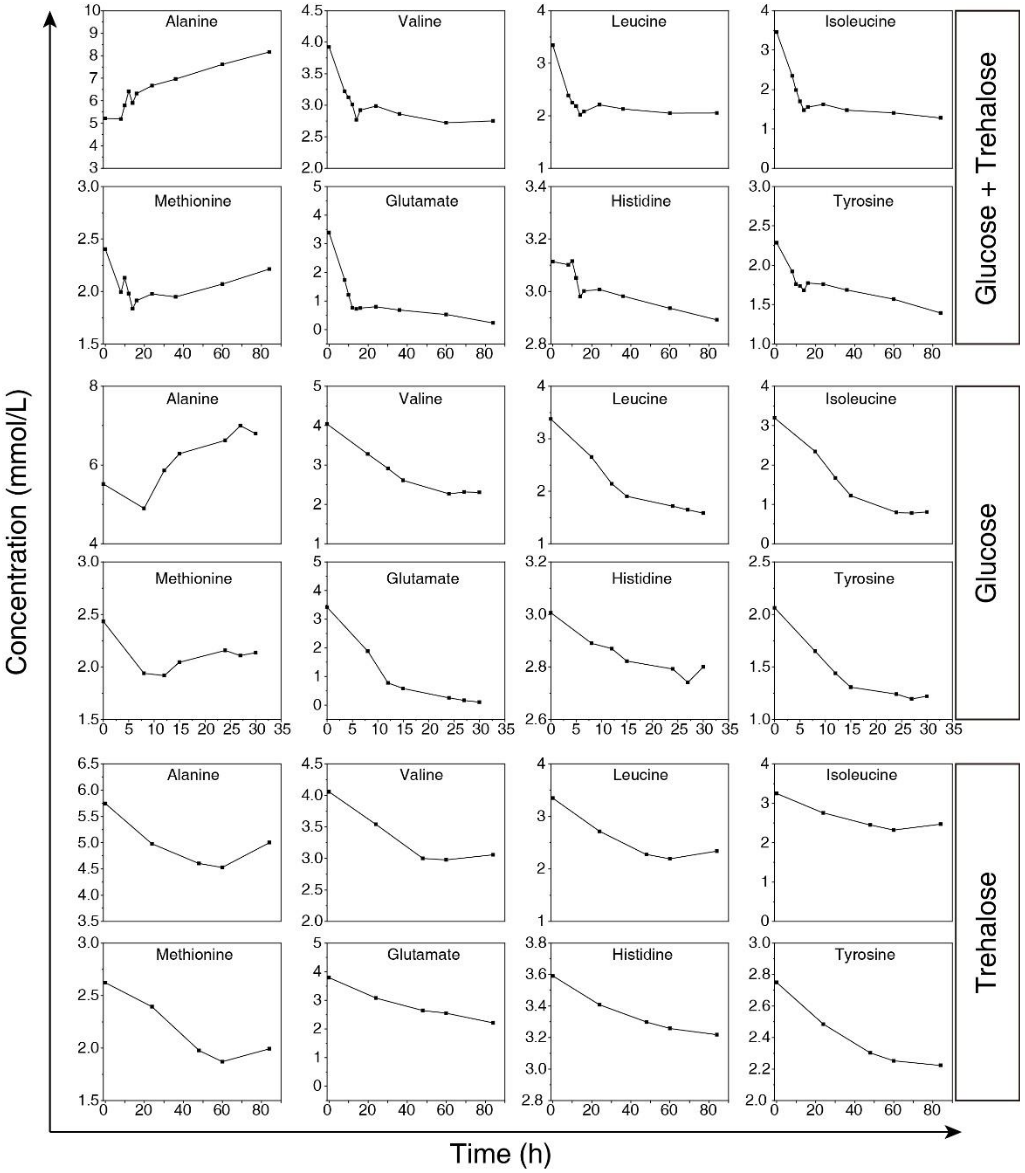

**Supplementary Fig. 2. Amino acid concentrations in the medium under different conditions.**

Under mixed carbon source conditions, amino acids were rapidly utilized along with the consumption of glucose. After glucose depletion, the uptake of amino acids slowed down, and they were not exhausted until the end of fermentation. Even under the sole glucose conditions, amino acids remained abundant.

### Supplementary Fig. 3

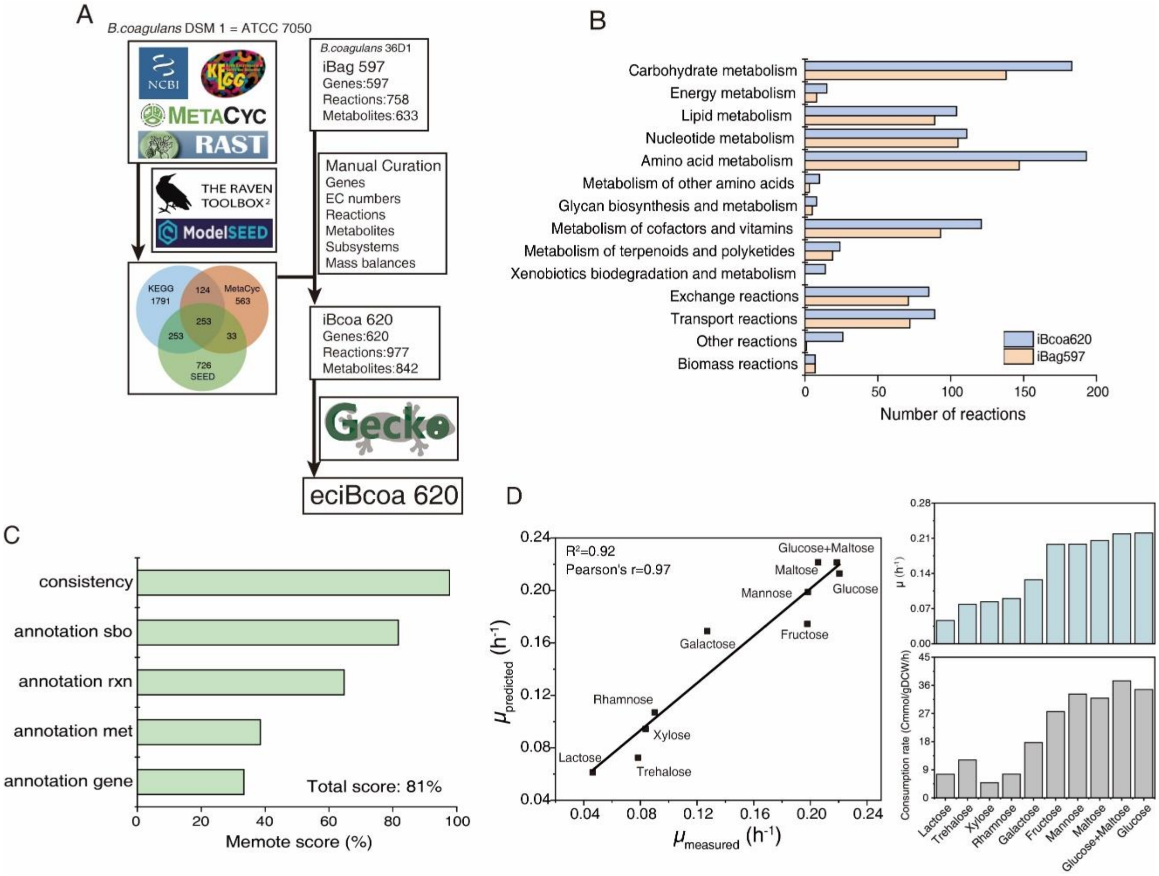

**Supplementary Fig. 3. Reconstruction and evaluation of genome-scale metabolic model.** (A) The construction process of the genome-scale metabolic model and enzyme-constrained model of *Bacillus coagulans* DSM 1 = ATCC 7050. The model iBcoa620 is primarily derived from two sources: a collection of coarse models from different databases and the existing model iBag597 of *B. coagulans* 36D1. The enzyme-constrained model was constructed using the latest GECKO toolbox. (B) In comparison to iBag597, iBcoa620 exhibits an increased number of reactions in most KEGG pathways, particularly in carbohydrate metabolism and amino acid metabolism. (C) The evaluation of iBcoa620 was conducted based on the online assessment system Memote, resulting in a final score of 81%, indicating a high-quality model. (D) The predictive capabilities of iBcoa620 were evaluated through shake flask experiments. Fermentation was carried out under different carbon source conditions (total carbon source of 40 g/L), and biomass and carbon source concentrations were determined during the culture process for the calculation of maximum specific growth rate and carbon source uptake rates. The measured sugar consumption rates were used to constrain the model to predict the maximum growth rates. The Pearson correlation coefficient between the experimentally measured values and the predicted values was 0.97.

### Supplementary Fig. 4

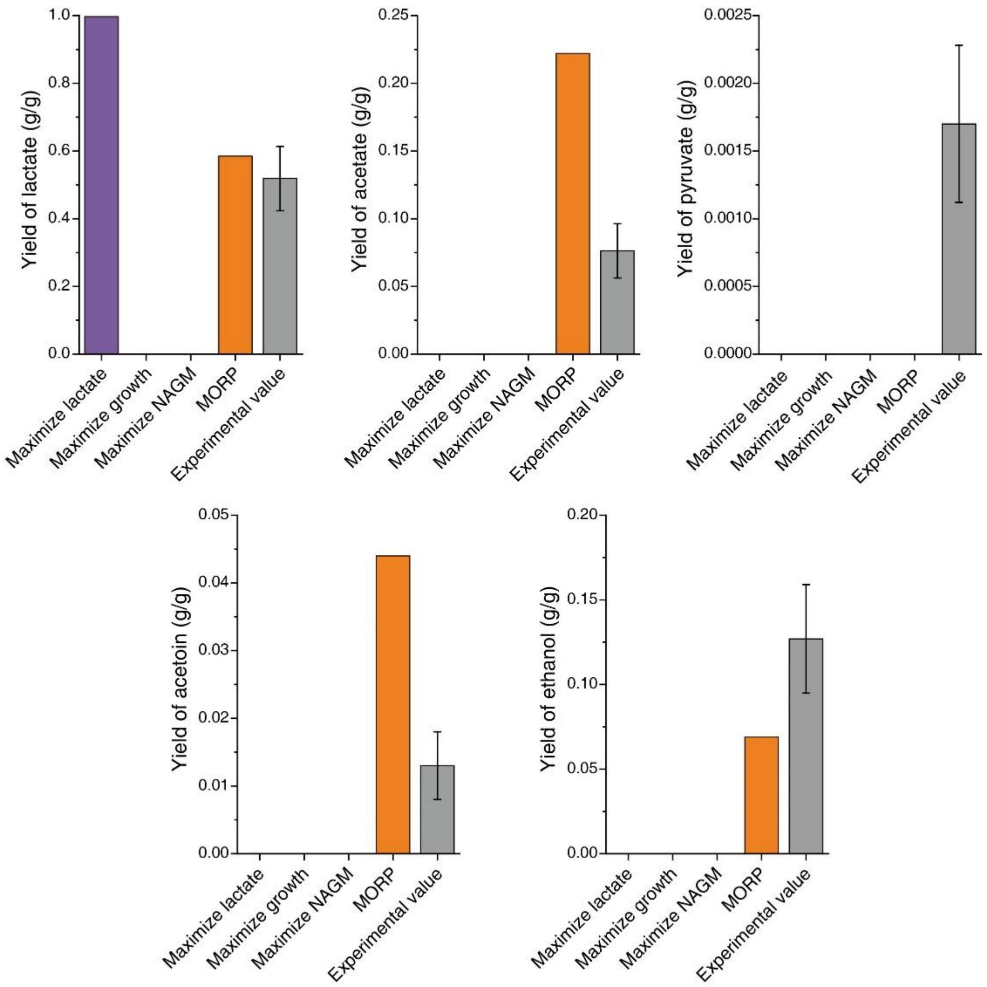

**Supplementary Fig. 4. Product yields in $P_{tre}$ of the different objective functions compared with the experimental results.**

The lactate yield of $P_{tre}$ predicted by MORP was consistent with the experimental value. MORP predicted the secretion of acetate and acetoin although the yields were higher than the experimental values. No objective function predicted pyruvate production, whose experimental yield was extremely low. MORP predicted ethanol secretion and the yield was closer to the experimental value.

### Supplementary Fig. 5

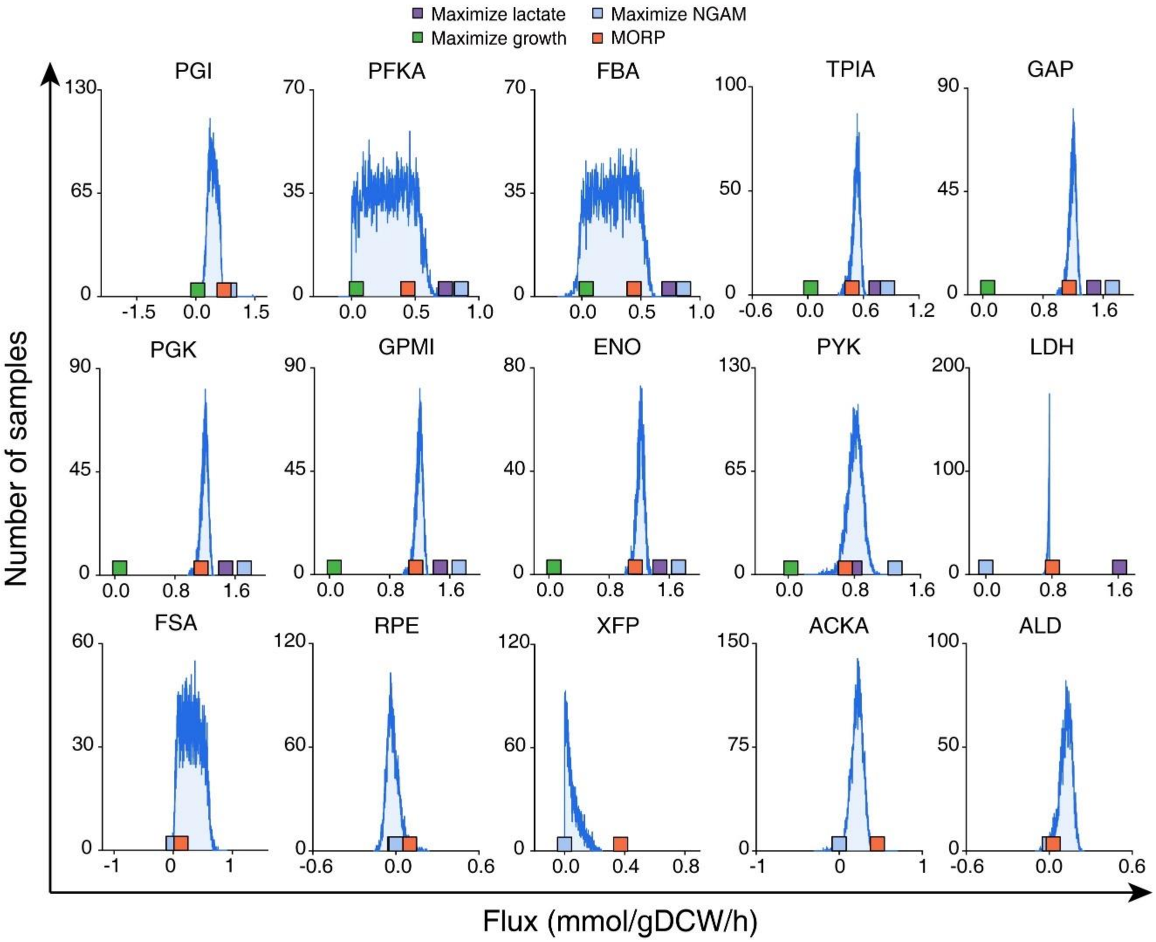

**Supplementary Fig. 5. Comparison between the sampled fluxes and the fluxes predicted by various objective functions for reactions in the central carbon metabolism.**

Random sampling was performed to obtain unbiased flux distributions, in which the rates of exchange metabolites including trehalose, organic acids and amino acids were constrained by the measured values of $P_{tre}$ 48 h. Within the solution space, 10,000 flux distributions were randomly sampled and fluxes of reactions in the central carbon metabolism are shown in the figure. The corresponding fluxes predicted using various objective functions are also plotted for comparison. The fluxes predicted by MORP were generally within the ranges of the sampled fluxes, while the fluxes by other objective functions deviated from the ranges.

### Supplementary Fig. 6

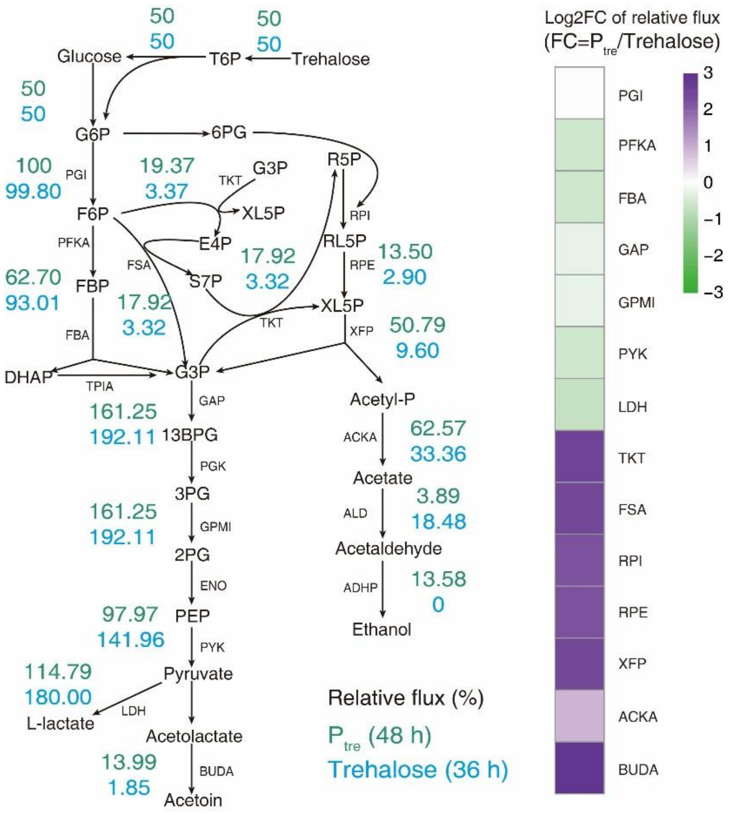

**Supplementary Fig. 6. Comparison of the relative flux and log2FC between $P_{tre}$ and sole trehalose.**

Under the sole trehalose condition, dFBA was used to simulate the utilization of trehalose, and the objective function was set to maximize growth. With the premise of consistent predicted phenotype and experimental results, the relative fluxes in central carbon metabolism of 36 h were compared with those of $P_{tre}$. The results confirmed that under the condition of sole trehalose, lactate was produced through the homolactic fermentation pathway. Additionally, $P_{tre}$ exhibited the largest variation in fluxes of the heterolactic fermentation pathway, further confirming the metabolic transition in $P_{tre}$. T6P, trehalose 6 - phosphate; G6P, glucose 6 - phosphate; F6P, fructose 6 - phosphate; FBP, fructose 1,6 - bisphosphate; DHAP, dihydroxyacetone phosphate; G3P, glyceraldehyde 3 - phosphate; 13BPG, glycerate 1,3 - bisphosphate; 3PG, 3-phosphoglycerate; 2PG, 2-Phospho-D-glycerate; PEP, phosphoenolpyruvate; 6PG, gluconate 6 - phosphate; E4P, erythrose 4 - phosphate; S7P, sedoheptulose 7-phosphate; R5P, ribose 5 - phosphate; RL5P, ribulose 5 - phosphate; XL5P, xylulose 5-phosphate; Acetyl-P, Acetyl phosphate; PGI, glucose-6-phosphate isomerase; PKFA, 6-phosphofructokinase; FBA, fructose-1,6-bisphosphate aldolase; GAP, glyceraldehyde-3-phosphate dehydrogenase; PGK, phosphoglycerate kinase; GPMI, phosphoglycerate mutase; ENO, enolase; PYK, pyruvate kinase; LDH, L-lactate dehydrogenase; RPI, ribose 5-phosphate isomerase; RPE, ribulose-phosphate 3-epimerase; XFP, phosphoketolase; FSA, fructose-6-phosphate aldolase; TKT, transketolase; BUDA, acetolactate decarboxylase; ACKA, acetate kinase; ALD, aldehyde dehydrogenase; ADHP, alcohol dehydrogenase.

### Supplementary Fig. 7

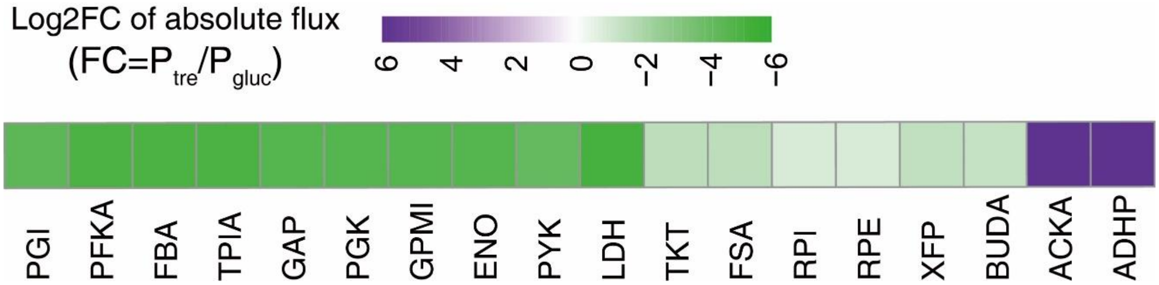

**Supplementary Fig. 7. Absolute flux changes of reactions in the central carbon metabolism pathway in $P_{tre}$ versus $P_{gluc}$.**

Absolute metabolic flux changes in the CCM in $P_{tre}$ versus $P_{gluc}$. In terms of absolute flux comparison, the flux variation in the phosphoketolase pathway (RPI, RPE, XFP) is minimal. This results in the allocation of a greater proportion of carbon source flow toward this pathway at lower trehalose uptake rate. Consequently, it leads to differences in the production of lactate and acetate. log2FC values were calculated using the mean of gene expression levels between the two conditions.

### Supplementary Fig. 8

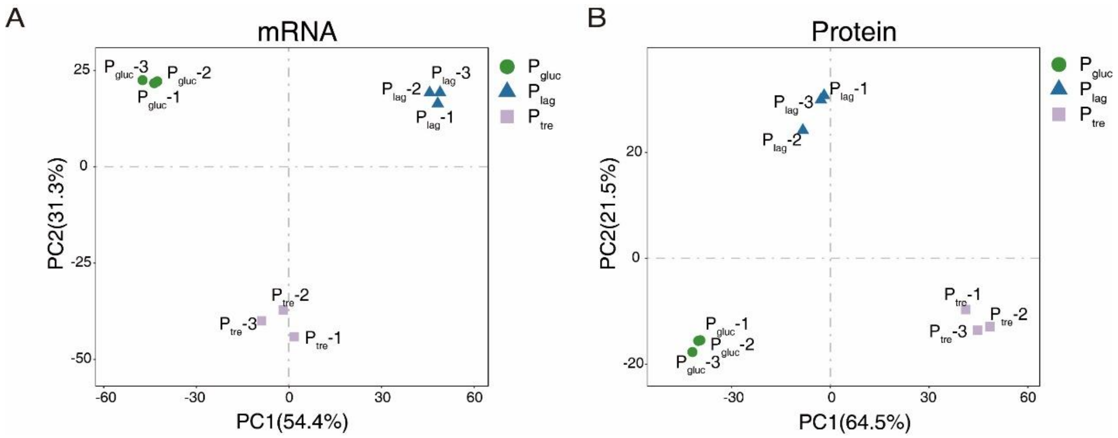

**Supplementary Fig. 8. Principal component analysis (PCA) plotting for three phases based on transcriptomic and proteomic data.**

PCA is a common method used to assess within-group repeatability and between-group differences. Closer proximity between two samples indicates higher similarity, while greater dispersion suggests significant differences between samples. Similarly, closer proximity between groups indicates smaller between-group differences, while distinct separation indicates significant between-group differences. PCA analyses were conducted separately for transcriptome (A) and proteome (B) data obtained at 7h, 14h, and 48h time points. The first two principal components (PC1, PC2), which accounted for 85% of the cumulative reliability, were plotted for the three groups. From the plot, it can be observed that the within-group samples tend to cluster together for both the transcriptome and proteome data, indicating high repeatability. Furthermore, the clear separation between the groups indicates noticeable overall differences between different phases.

### Supplementary Fig. 9

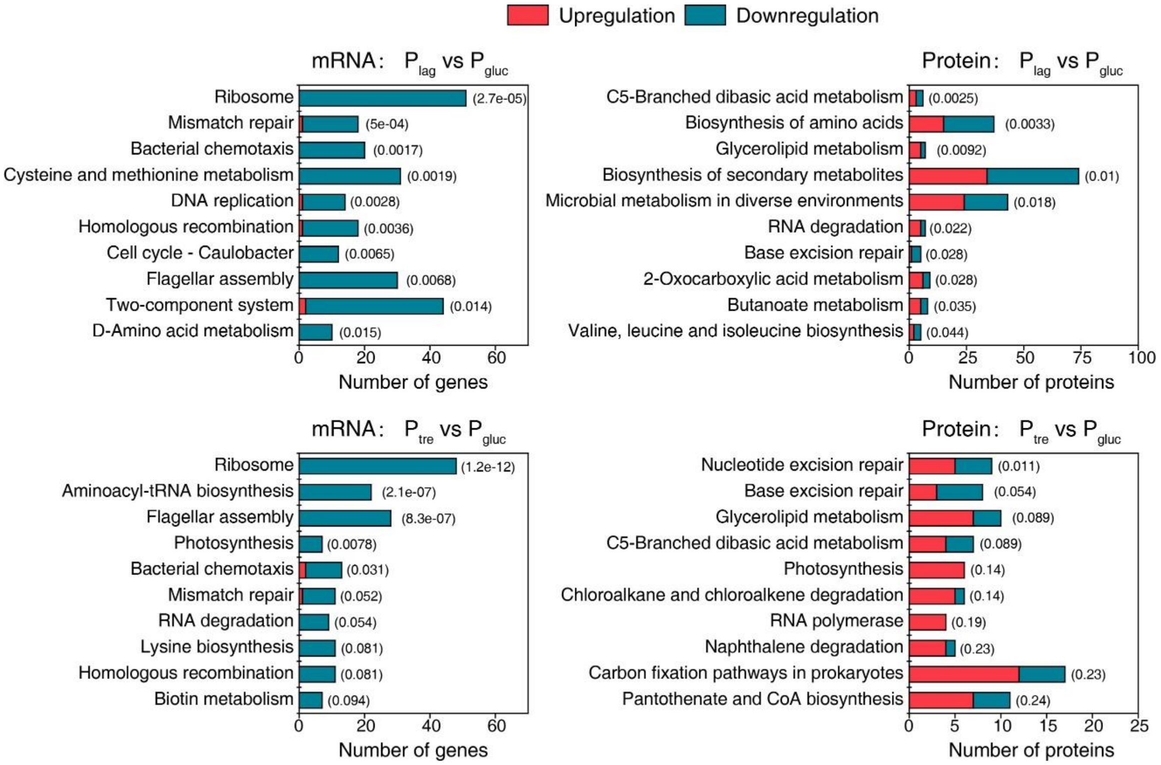

**Supplementary Fig. 9. KEGG enrichment analysis of differentially expressed transcripts and proteins between different phases.**

Number in parentheses represents the *P* value of the enrichment.

### Supplementary Fig. 10

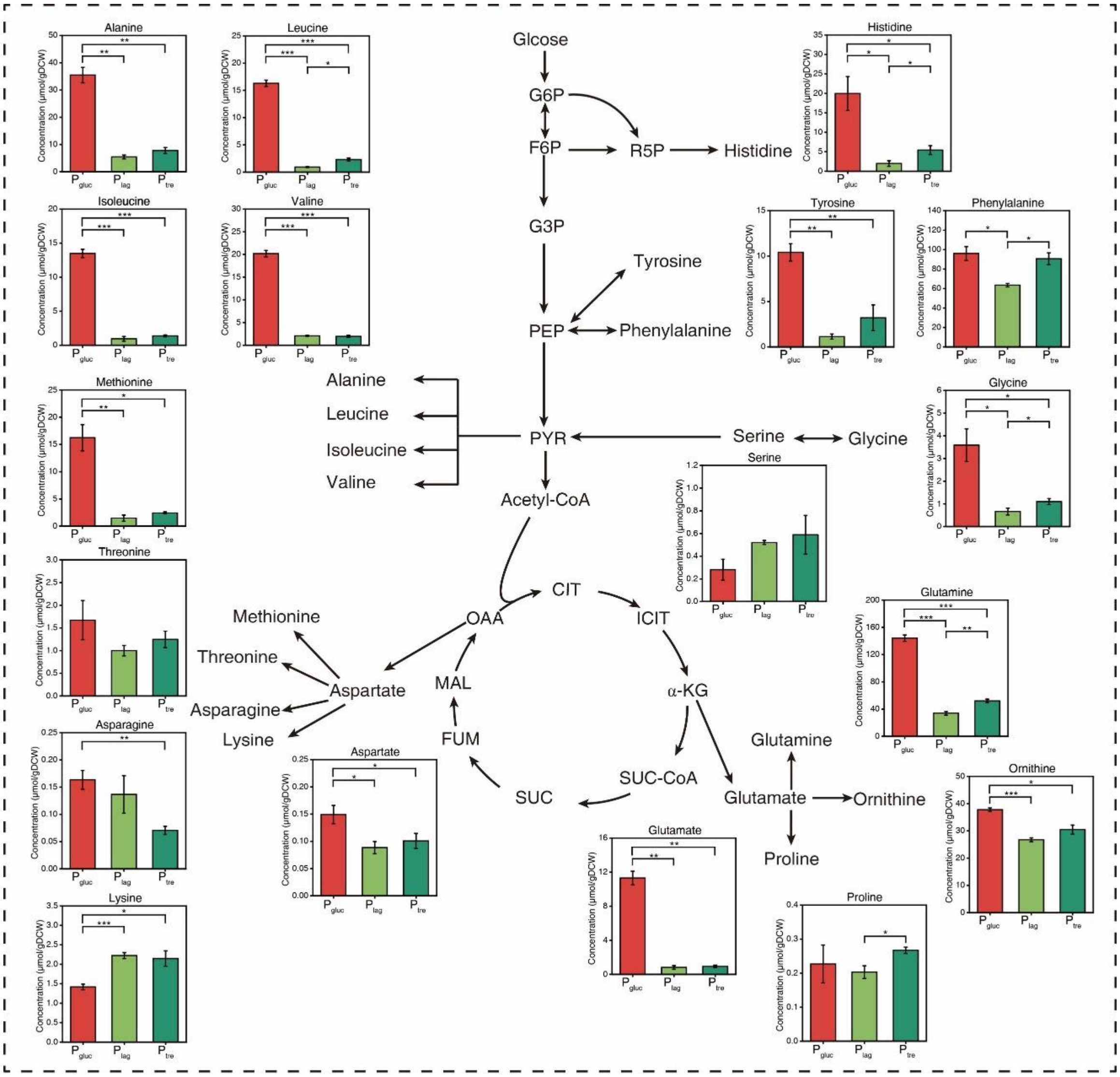

**Supplementary Fig. 10. Concentrations of intracellular amino acids of different phases.** Data are mean ± s.d. of three biological replicates. Statistical analysis was performed using two-sided *t*-test. Statistically significant differences are described as follows: *** $P$<0.001, ** $P$<0.01, * $P$<0.05.

### Supplementary Fig. 11

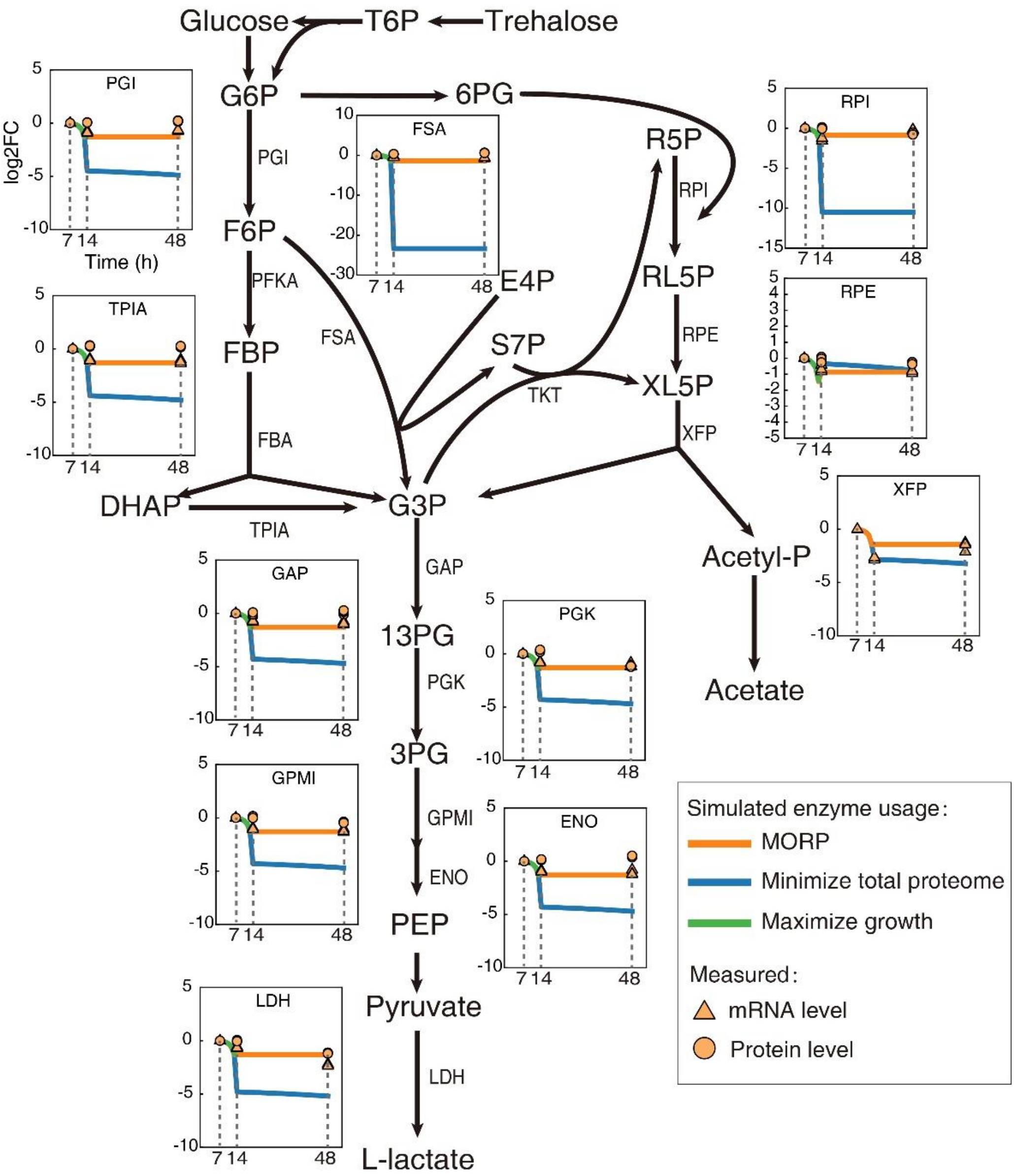

**Supplementary Fig. 11. Comparison between the simulations and the omics data for the CCM reactions .**

Comparison of enzyme usage simulated by dMORP and the objective function of minimizing the total proteome with omics data. Taking the PGI reaction as an example, the log2FC value represents the change in enzyme usage, mRNA or protein level relative to the 7 h point. The omics data points are all three biological replicates.

**Supplementary Fig. 12**

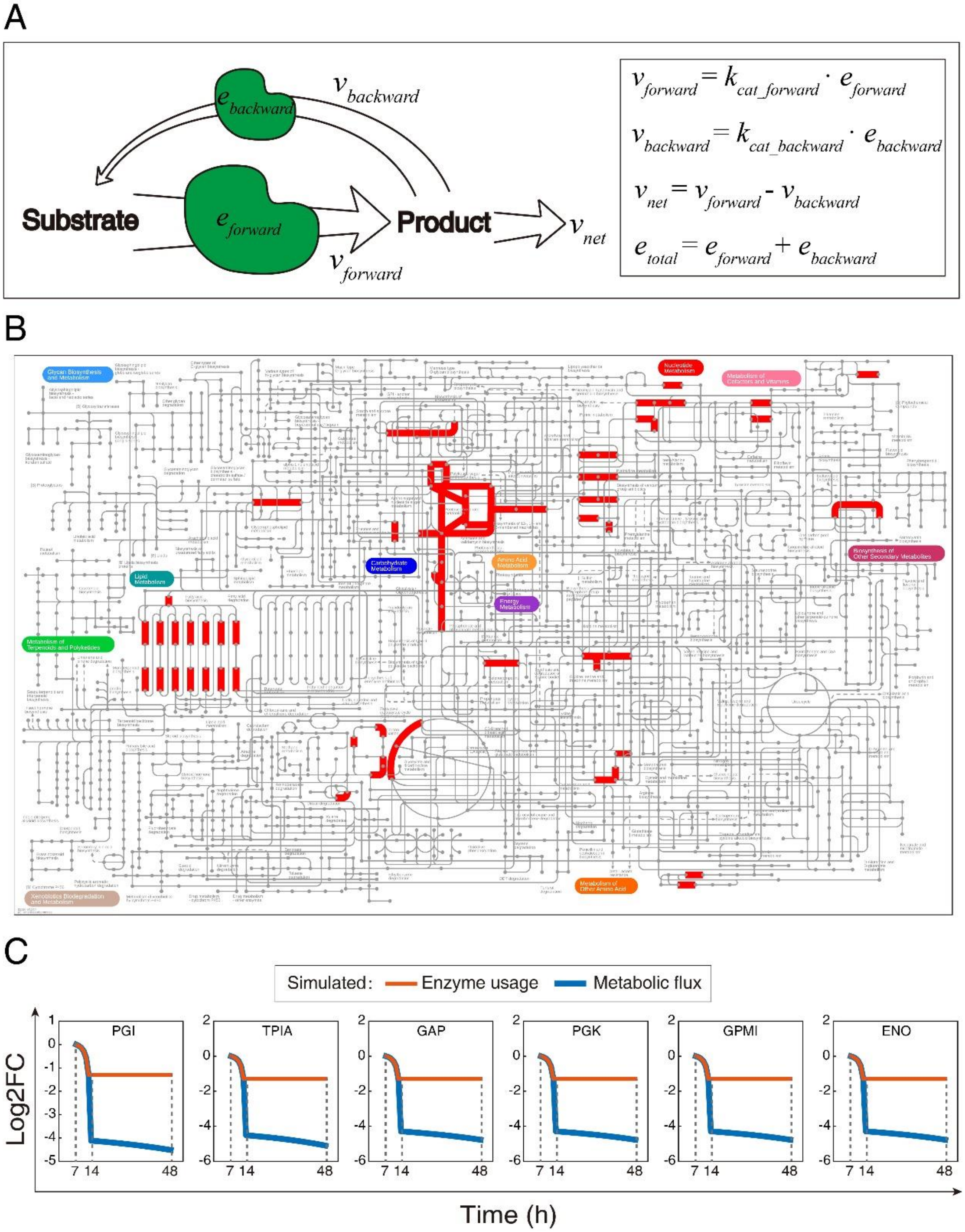

**Supplementary Fig. 12. Predicted mismatch between enzyme usage and metabolic flux.**

(A) MORP predicts the mismatch between enzyme usage and metabolic flux with enzyme-constrained models. (B) The reactions (red lines) with mismatch between enzyme usage and metabolic flux. (C) Comparison of the changes of glycolytic reactions between the enzyme usage and metabolic flux. dMORP was used to simulate for the period from 14 to 48h. The log2FC value represents the change in enzyme usage and metabolic flux relative to the 7 h point.

### Supplementary Fig. 13

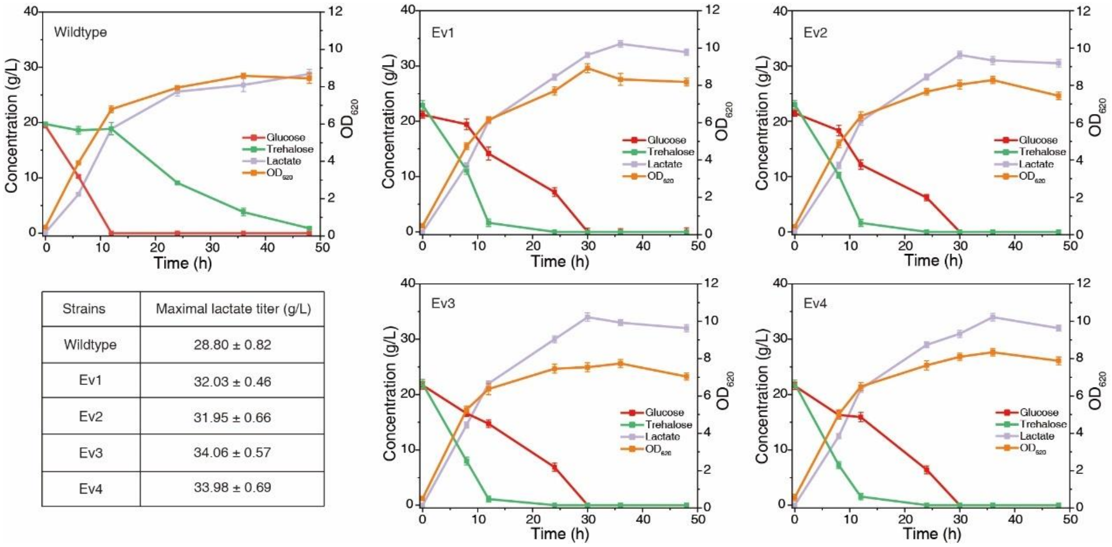

| Strains | Maximal lactate titer (g/L) |
|---|---|
| Wildtype | 28.80 ± 0.82 |
| Ev1 | 32.03 ± 0.46 |
| Ev2 | 31.95 ± 0.66 |
| Ev3 | 34.06 ± 0.57 |
| Ev4 | 33.98 ± 0.69 |

**Supplementary Fig. 13. Comparison of fermentation phenotypes of wildtype strain and evolved strains with mixed carbon sources in shake flasks.**

The wildtype strain exhibited a clear hierarchical utilization phenomenon and residual trehalose even can be detected after 48 h. In contrast, the four evolved strains (Ev1, Ev2, Ev3, Ev4) demonstrated the ability to simultaneously utilize glucose and trehalose effectively. The uptake rate of trehalose was higher than that of glucose, and both carbon sources were almost depleted after 30 h. In terms of maximum lactate production, the evolved strains also exhibited higher lactate production capacity. Among the four evolved strains, Ev3, which showed higher lactate production, was selected for subsequent 5L bioreactor experiments. Data are mean ± s.d. of three biological replicates.

**Supplementary Fig. 14**

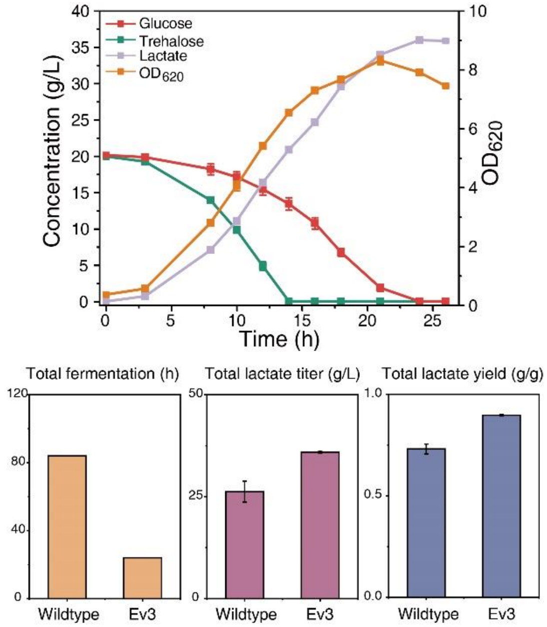

**Supplementary Fig. 14. Fermentation on the mixture of glucose and trehalose in 5 L bioreactor by the evolved strain Ev3.**

Ev3 demonstrated simultaneous utilization of glucose and trehalose, greatly shortened fermentation cycle and improved total lactate yield. Data are shown with mean ± s.d. of three biological replicates.

### Supplementary Table 1

**Supplementary Table 1. Physiological parameters of *B. coagulans* under different carbon sources.**

$P_{gluc}$ and $P_{tre}$ represent the glucose and trehalose consumption phase under mixed carbon sources, respectively. Ev3 represents the evolved strain that can co-utilize glucose and trehalose. Data in table are mean ± s.d. of three biological replicates.

| | ATCC7050 | | | | Ev3 (glucose+trehalose) |
|---|---|---|---|---|---|
| | $P_{gluc}$ | $P_{tre}$ | Glucose | Trehalose | |
| $\mu_{max}$ (/h) | 0.313±0.008 | / | 0.300±0.002 | 0.053±0.004 | 0.258±0.002 |
| $q_s$ (g/gDCW/h) | 2.118±0.177 | 0.107±0.039 | 1.748±0.020 | 0.423±0.027 | 1.566±0.046 |
| $V_S$ (g/L/h) | 1.759±0.061 | 0.217±0.053 | 1.333±0.007 | 0.218±0.003 | 1.671±0.002 |
| $V_{lactate}$ (g/L/h) | 1.534±0.078 | 0.114±0.045 | 1.175±0.009 | 0.185±0.012 | 1.499±0.007 |
| $Y_{X/S}$ (g/g) | 0.119±0.007 | / | 0.075±0.001 | 0.039±0002 | 0.062±0.001 |
| $Y_{lactate/S}$ (g/g) | 0.898±0.009 | 0.526±0.041 | 0.882±0.006 | 0.851±0.044 | 0.895±0.004 |
| $Y_{citrate/S}$ (g/g) | 0.007±0.003 | / | 0.002±0.001 | / | 0.003±0.001 |
| $Y_{pyruvate/S}$ (g/g) | 0.001±0.000 | 0.002±0.001 | 0.001±0.000 | 0.001±0.000 | 0.001±0.000 |
| $Y_{acetate/S}$ (g/g) | 0.025±0.001 | 0.100±0.039 | 0.024±0.002 | 0.069±0.005 | 0.025±0.001 |
| $Y_{acetoin/S}$ (g/g) | 0.009±0.002 | 0.031±0.012 | 0.024±0.001 | 0.009±0.000 | 0.030±0.002 |
| $Y_{CO2/S}$ (g/g) | 0.012±0.000 | 0.038±0.002 | 0.013±0.000 | 0.022±0.001 | 0.010±0.000 |
| $Y_{ethanol/S}$ (g/g) | 0.008±0.001 | 0.127±0.032 | 0.007±0.000 | 0.010±0.000 | 0.009±0.000 |
| Carbon balances (%) | 104.952+1.752 | 83.277±2.008 | 104.022±1.184 | 95.257±2.481 | 102.681±0.164 |

### Supplementary Table 2

**Supplementary Table 2. Amino acid sequence changes of *crr*.**

Four evolved strains exhibit identical mutation in the *crr* gene, resulting in amino acid sequences listed in the table.

| Gene | Original sequence（160aa） | Mutated sequence（49aa） |
|---|---|---|
| *crr*<br>BF29_RS12935 | MFKLFQKKPKEERIFAPVRGQVVAIEEVPDP<br>TFSEKMMGDGIAVKPTEGLLVAPFDGEVVQ<br>VFPTKHAIGLRGASGLELLIHIGLETVTLNRE<br>GFETAVKDGDKVKKGDTLITFDLEWIQEKAA<br>DSITSIVVTNGEAVHLDKKTSLDAVPGETVV<br>MTAQL | MYIDLEWIQEKAADSITSIVV<br>TNGEAVHLDKKTSLDAVPG<br>ETVVMTAQL |
